# Crystal or Jelly? Effect of Color on the Perception of Translucent Materials with Photographs of Real-world Objects

**DOI:** 10.1101/2021.10.18.464695

**Authors:** Chenxi Liao, Masataka Sawayama, Bei Xiao

## Abstract

Translucent materials are ubiquitous in nature (e.g. teeth, food, wax), but our understanding of translucency perception is limited. Previous work in translucency perception has mainly used monochromatic rendered images as stimuli, which are restricted by their diversity and realism. Here, we measure translucency perception with photographs of real-world objects. Specifically, we use three behavior tasks: binary classification of “translucent” versus “opaque”, semantic attribute rating of perceptual qualities (see-throughness, glossiness, softness, glow and density), and material categorization. Two different groups of observers finish the three tasks with color or grayscale images. We find that observers’ agreements depend on the physical material properties of the objects such that translucent materials generate more inter-observer disagreements. Further, there are more disagreements among observers in the grayscale condition in comparison to that in color condition. We also discover that converting images to grayscale substantially affects the distributions of attribute ratings for some images. Furthermore, ratings of see-throughness, glossiness, and glow could predict individual observers’ binary classification of images in both grayscale and color conditions. Lastly, converting images to grayscale alters the perceived material categories for some images such that observers tend to misjudge images of food as non-food and vice versa. Our result demonstrates color is informative about material property estimation and recognition. Meanwhile, our analysis shows mid-level semantic estimation of material attributes might be closely related to high-level material recognition. We also discuss individual differences in our results and highlight the importance of such consideration in material perception.

## Introduction

Many materials in our daily life are translucent, which allow some of light to penetrate the object, refract, and scatter multiple times throughout the body of the medium, before exiting from a different location on the surface. Translucent materials are ubiquitous in life including skin, teeth, fruits, liquid, crystals, glass, plastics, and wax. Yet, in comparison to opaque materials, relatively little is understood about translucent appearance (Fleming,2017;Gigilashvili, Thomas, Hardeberg, & Pedersen,2021). Translucency is challenging to study due to several reasons. First, the physical process of translucency is complex, involving surface reflection and sub-surface scattering (see (Gkioulekas et al.,2013;Xiao et al.,2014) for detailed description of physical model of sub-surface scattering and translucency perception). Previous work shows that physical material properties, lighting, shape, and context all affect the appearance of translucent objects (Fleming & Bülthoff,2005;Motoyoshi,2010;Nagai et al.,2013;Xiao et al.,2014;Marlow, Kim, & Anderson,2017;Chowdhury, Marlow, & Kim,2017;Sawayama et al.,2019;Tamura, Higashi, & Nakauchi,2018;Xiao, Zhao, Gkioulekas, Bi, & Bala,2020;Marlow & Anderson,2021;Gigilashvili, Shi, et al.,2021;Gigilashvili, Thomas, Hardeberg, & Pedersen,2021), and it still remains unknown how humans extract intrinsic translucent material properties from images. Second, there are many different kinds of translucent materials in real-life. Different types of materials (e.g. skin versus wax) have different physical generative processes. Even though these materials can be all described as “translucent”, humans have no trouble visually discriminating them.

Humans can often report whether a surface is glossy or matte with verbal rating, even though perception of gloss is multidimensional (Ferwerda, Pellacini, & Greenberg,2001). But translucent appearance is difficult to be verbally described and is likely to be high dimensional. Using rendered images, most previous works have measured translucent appearance using a single task (e.g. asking observers to rate the apparent translucency, asking observers to judge which of the two images seems to be more translucent, or matching the images based on the perceived translucency by using a slider to adjust one particular physical parameter) (Motoyoshi,2010;Nagai et al.,2013;Xiao et al.,2020). Such measurements may not fully reflect how humans perceive and recognize translucent materials in the real world. In this paper, we measure material perception using photographs of real-world objects in three different tasks, namely, binary classification of translucency (Experiment 1), semantic attributes rating (Experiment 2), and material categorization (Experiment 3) with the goal of understanding translucency perception in the wild with a focus on the role of color.

### Previous findings on translucency

Previous research has focused on exploring the relationship between physical parameters in the sub-surface scattering and translucent appearance using rendered images. Fleming and Bülthoff have studied the effect of direction of illumination on translucency perception of rendered objects with relatively simple rendering models and geometry (Fleming & Bülthoff,2005). One of their findings is that observers seem to be poor at “discounting” the effects of light source direction, and translucent objects tend to appear more translucent when they are illuminated from behind. Xiao et al. use a more complex 3D shape, a “Lucy” model, rendered with more sophisticated models, and show that the lighting direction has a strong effect on translucent appearance with non-isotropic scattering phase functions (Xiao et al.,2014). Gkioulekas et al. find translucent edges have specific qualitative features as the result of combination of scattering, refraction, and reflection, and these features can distinguish translucent edges from edges in opaque objects (Gkioulekas, Walter, Adelson, Bala, & Zickler,2015). Using rendered images, previous work has also proposed image cues for perceiving translucency (see a recent review (Gigilashvili, Thomas, Hardeberg, & Pedersen,2021) for details). Specifically, Motoyoshi et al. show that the “relationship with respect to contrast and sharpness between the specular highlights and the non-specular body” is a robust cue for perceived translucency (Motoyoshi,2010). Nagai et al. find that perceptual translucency depends on local image features, such as mean luminance, within specific image regions (Nagai et al.,2013). Even though these works provide testable image cues for translucency perception, it is not clear whether they can be generalized to real-world materials beyond rendered images.

### Role of color in translucency

Nearly all previous work in translucency perception has used monochromatic images (Motoyoshi,2010;Nagai et al.,2013;Xiao et al.,2020). The use of monochromatic images in translucency perception is largely due to the lack of high-quality renderings of physically-plausible colored images of translucent appearance. Even though there are significant improvements in spectral rendering of translucent materials such as wax, jade, and skin (Jimenez, Whelan, Sundstedt, & Gutierrez,2010;Brunton, Arikan, Tanksale, & Urban,2018), these rendering methods may not extend to realistically replicate the variety of translucent appearances in real life, such as food. Some recent studies examine translucency in a more realistic setting: Gigilashvili et al. have used real physical objects (an artwork collection Plastique made of resin) to explore potential appearance ordering system that implies translucency (Gigilashvili, Thomas, Pedersen, & Hardeberg,2021); Chadwick et al. used color photographs of translucent liquids (glasses of milky tea) when they compared perceptual performance with real and computer-generated stimuli (A. C. Chadwick, Cox, Smithson, & Kentridge,2018). In reality, color could play important roles in translucent appearance as shown in Figure 1. For example, some low-level color-related image cues (e.g. saturation) might affect perceived translucency (e.g. the color gradients in yellow microcrystalline wax cube is an important cue). On the other hand, removing color might also alter the state of the object through association (e.g. cooked shrimp might appear to be uncooked when the image is shown in grayscale), which in turn affects the observer’s material perception (e.g. uncooked shrimp is usually perceived to be more translucent). Previous works find color interacts with perception of surface gloss (Xiao & Brainard,2008;Nishida, Motoyoshi, & Maruya,2011), perception of fabrics (Xiao, Bi, Jia, Wei, & Adelson,2016;Toscani, Milojevic, Fleming, & Gegenfurtner,2020), perception of transparent objects (D’Zmura, Colantoni, Knoblauch, & Laget,1997;Ennis & Doerschner,2021), and perception of object states such as wet or bleached (Sawayama, Adelson, & Nishida,2017;Okawa et al.,2019;Toscani et al.,2020). On the other hand, a previous study on material classification from photographs using the Flickr Material Dataset finds removing color does not significantly affect classification accuracy (Sharan, Rosenholtz, & Adelson,2014). However, little is known about how removing color affects the perception of more diverse materials including translucent objects. Some studies (Fleming & Bülthoff,2005;A. C. Chadwick et al.,2018) have pointed out that saturation variations can affect the perceived translucent appearance, but they are insufficient on their own to produce an impression of translucency. It is also not clear whether color affects material appearance mostly through low-level processing of color-related image cues or through high-level association with object identity or recognition memory (Wichmann, Sharpe, & Gegenfurtner,2002). A previous study has also studied translucency perception of a colour blind observer, suggesting that some aspects of translucence perception do not depend on regions critical for color and texture processing (A. Chadwick, Heywood, Smithson, & Kentridge,2019). In this paper, by systematically measuring the effect of color across different tasks, we aim to discover the relationship between different hierarchical processes involved in material perception.

**Figure 1:**
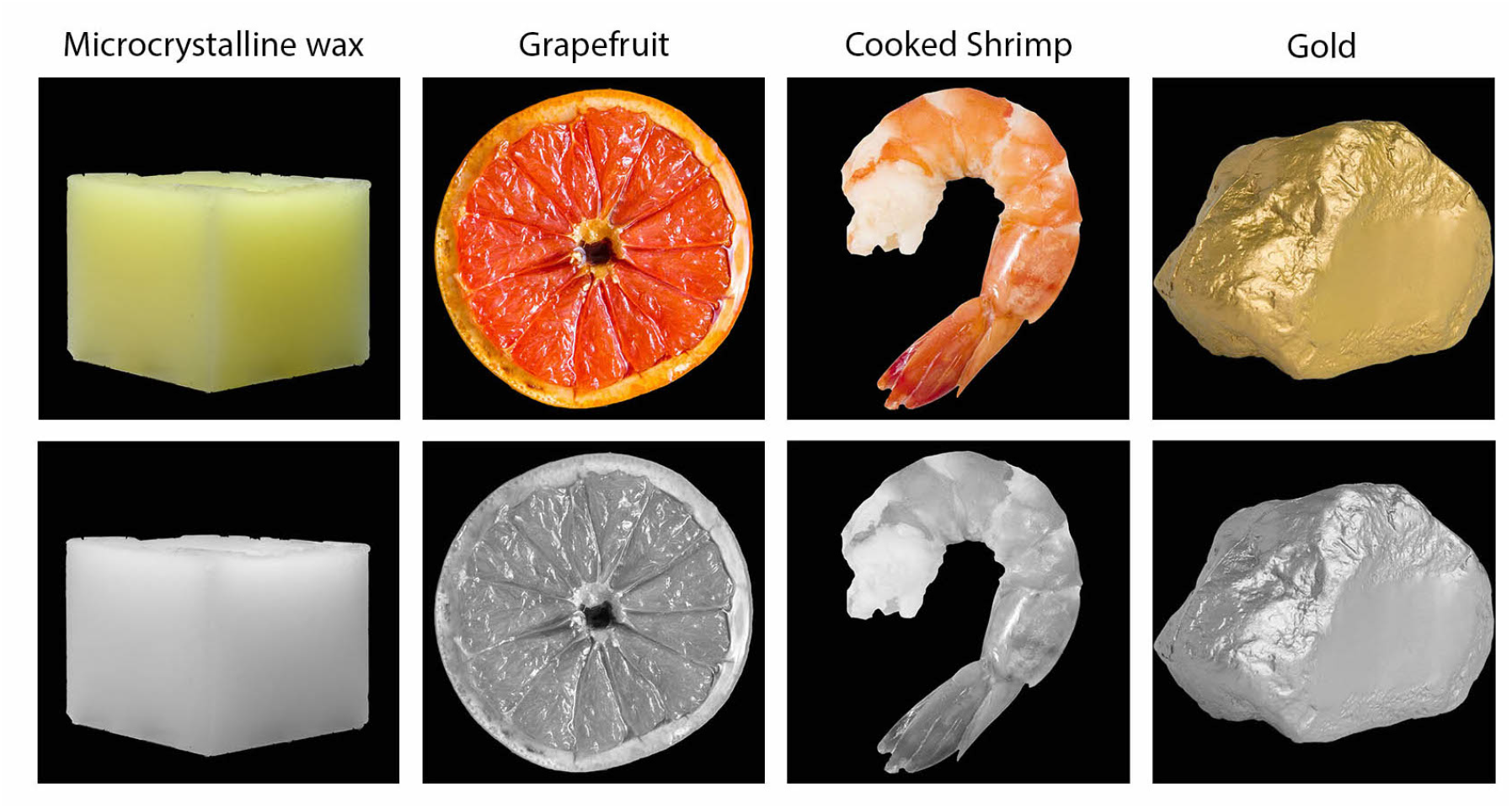
Illustration of how converting images to grayscale affects various aspects of material perception of translucent and opaque objects, including low-level image cues, mid-level estimation of material attributes, and high-level material recognition. From left to right, the microcrystalline wax (parawax) cube appears less translucent and less “see-through” in grayscale than in color; the grapefruit looks less juicy and might be recognized as a fake fruit in grayscale; the cooked shrimp might look uncooked in grayscale; the chunk of gold might appear plastic in grayscale.

### High level material perception and individual difference

Relatively few studies have examined high-level semantic material perception such as assigning semantic class of materials from real-world images or paintings (Sharan et al.,2014;Sharan, Rosenholtz, & Adelson,2009;Fleming, Wiebel, & Gegenfurtner,2013;Baumgartner, Wiebel, & Gegenfurtner,2013;Zuijlen, Pont, & Wijntjes, 2020;Wijntjes, Spoiala, & De Ridder,2020;Di Cicco, Wijntjes, & Pont,2020). Specifically, Sharan et al. find that humans can recognize materials in rapid visual presentation within pre-designed categories (Sharan et al.,2009). Fleming et al. find that ratings of subjective qualities (e.g. glossiness, colourfulness, roughness) could account for class membership with 90% accuracy (Fleming,2014). Fleming also suggests that material estimation might be affected by high-level material recognition (Fleming,2017).

We assume that measuring material perception using photographs of real objects, instead of rendered images, might result in larger individual difference. Chadwick et al. have observed that the models explaining the variation in the psychophysical data differ among individuals (A. C. Chadwick et al.,2018). A recent investigation suggests that young children rely more heavily on small-scale local image features for material perception than older children and adults do (Balas, Auen, Thrash, & Lammers,2020). Individual difference has been found in color constancy (Lafer-Sousa, Hermann, & Conway,2015;Toscani, Gegenfurtner, & Doerschner,2017;Witzel, O’Regan, & Hansmann-Roth,2017;Emery & Webster,2019;Aston & Hurlbert,2017), and in color naming and categorization, mostly due to cultural and environmental factors (Webster,2015), but it has not been systematically quantified in material perception. In a recent work, Gigilashvili et al. discussed multiple challenges observers have faced due to the ambiguity of the concept of perceptual translucency such as the vague definition of “translucency” in terms of perception and the limited knowledge on how to quantify translucency (Gigilashvili, Thomas, Hardeberg, & Pedersen,2020). In this study, we analyze both pooled results across observers and individual data, and discuss individual difference which might arise in the results.

### Main questions and study overview

In this paper, we aim at examining the role of color in perceiving translucent appearance across several tasks using photographs of real-world objects. In addition, we aim to understand how these tasks are related both at the group and individual levels. Our main hypothesis is that material perception involves both high-level recognition and mid-level estimation of material attributes, and we hypothesize that color affects both processes. We will ask the following questions: Will converting images to grayscale change an object’s appearance from “translucent” to “opaque”? How does color affect the estimation of material-related attributes? Can removing color alter the recognition of material categories (e.g. from crystal to jelly)?

To answer these questions, we design three experiments. We use a between-group design in which different groups of observers perform the same tasks with color and grayscale images, respectively. In Experiment 1, two groups of observers perform binary classification judging whether the material of the object is “translucent” or “opaque” in either color or grayscale condition (see Figure 2 (II)). In Experiment 2, we measure semantic attribute ratings of the same images in color and grayscale. Specifically, observers rate five attributes using a 6-point Likert scale (see Figure 2 (III)). In Experiment 3, we use a material categorization task to reveal how observers recognize the material categories (see Figure 2 (IV)). Across the three experiments, we aim to reveal how color affects the estimation of material properties and material categorization by analyzing both between-group effects and individuals’ performances. In the mean time, we investigate to what degree these tasks are related, and whether we can use the results from semantic attribute ratings to characterize translucency classification.

**Figure 2:**
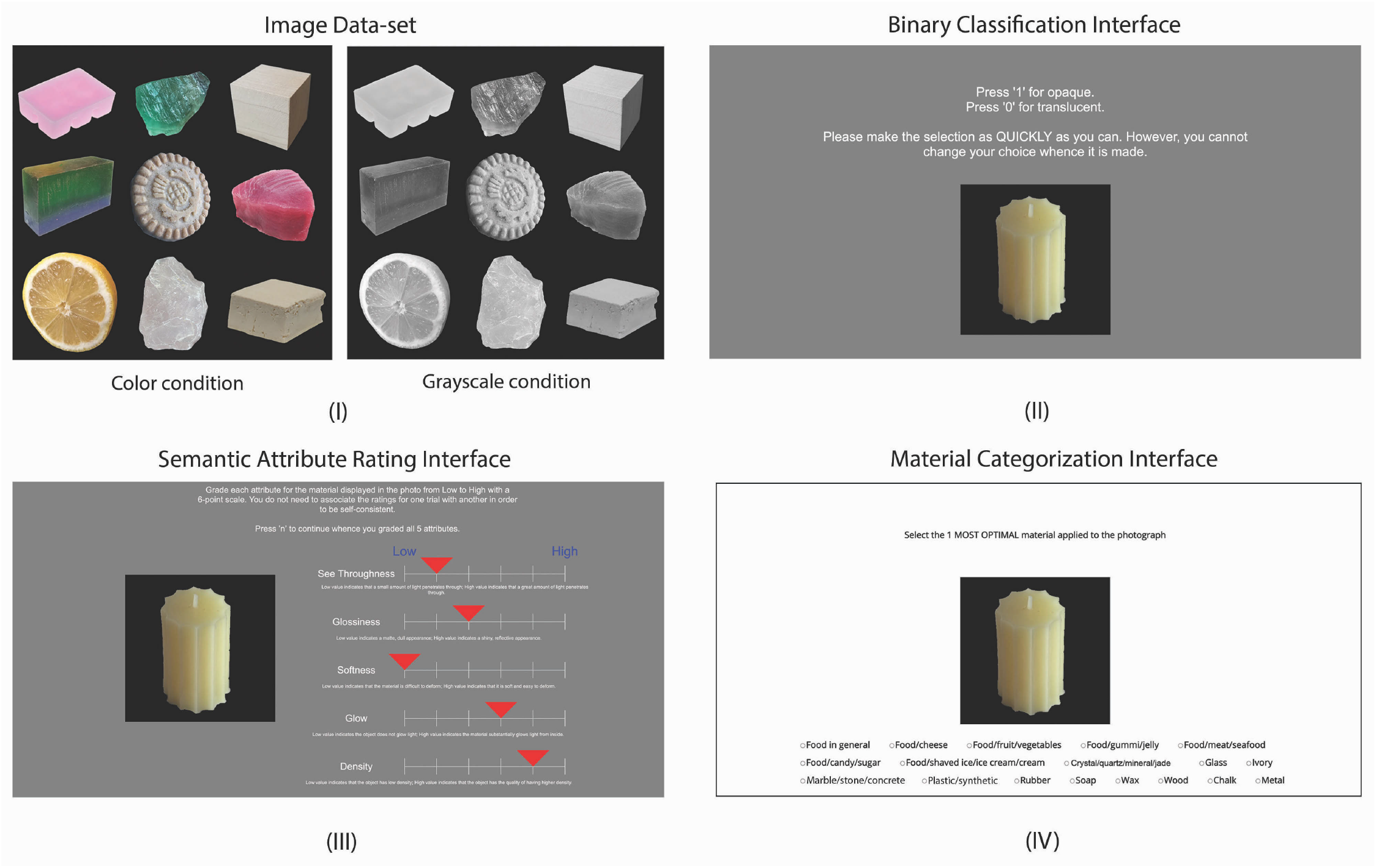
Stimuli and tasks. (I) Example stimuli in color and grayscale. (II) Binary classification experiment interface. Observers were asked to classify the material of the object as either “translucent” or “opaque”. (III) Semantic attribute rating experiment interface. Observers were asked to rate 5 translucency-related material attributes using a 6-point scale. 6 means high and 1 means low. (IV) Material categorization experiment interface. Observers were asked to choose the material category that best suits the object in the image among 18 categories.

## General method

### Image data-set and stimuli presentation

Our image data-set contains 300 photographs of everyday objects including fruit, meat, seafood, crystals, soap, wax, stone, plastics, etc., as experimental stimuli. The data-set consists of images downloaded from Google Images (with creative common licences) and photographs taken by the authors. Each photograph contains one object placed against a uniform black background, which is achieved by using a binary mask to filter the contour of the object before copying and pasting it onto an blank image. Even though, we try to make the ratio of the size of the object to the background consistent across the images, we do not control the lighting, image acquisition processes, and photography styles. Different from previous studies, we intend to sample the images from different sources in order have a representation of the richness of everyday visual experience.

The images from the data-set were presented to the observers in two separated conditions, color and grayscale, as shown in Figure 2 (I). In the color condition, the images were displayed in the original RGB color space. We created the grayscale images by removing the color from the RGB images. There are many methods to convert color images to grayscale. In the current study, our method was based on the lightness dimension in the CIELab color space. We assumed D65 as the reference to convert the sRGB images to grayscale. Here, we used the OpenCV function COLOR BGR2Lab to convert the BGR image into CIELab space, and then extracted, duplicated and concatenated the lightness dimension.

Due to the Covid-19 pandemic, the experiments were conducted online. Different from crowd-sourcing, we recruited the observers from American University. The experiments were built by PsychoPy and jsPsych and hosted on the online experiment platform Pavlovia.org (Peirce,2007;De Leeuw,2015). The observers had access to the experiment on their own computers and were instructed to use a device with at least 13-inch display and make the brightness of the screen at maximum level. The images were resized to 0.45 of the screen size used by the observers.

In both experiment, 60% of observers used laptop with MacOS system with 13-inch to 16-inch displays, and 40% used Windows system with 13-inch to 15-inch displays except 2 people used 24-inch displays.

### Observers and Experimental paradigm

Using a between-group design, we have one group of observers performing the tasks with color images, and a different group completing the tasks with grayscale images. In each condition, 20 observers first completed the binary classification, hence finished the semantic attribute rating experiment, and 15 of these observers also performed material categorization experiment. All observers are students from American University. The observers participated in the color experiments have a median age of 24. Observers participated in the grayscale experiments have a median age of 20. Overall, there are 12 females and 8 males in each group. Based on self-report, all observers speak fluent English though their native languages are different. All observers reported to have normal or corrected to normal visual acuity and have normal color vision. For all experiments, informed consent was obtained before experimentation. The procedures were conducted in accordance with the Declaration of Helsinki and were approved by the Human Research Ethics Advisory Panel at American University. We do not assume any prior difference between the two groups of observers in terms of life experiences with materials, special skills, and knowledge of image processing.

Figure 2 shows example stimuli and user interfaces for three experiments. The details of tasks and user interfaces are described in the subsequent experiment sections.

### Control experiment

We also conducted an experiment by using a different method of converting the color images to grayscale to test the extent to which the results are affected. We generated the grayscale images using the Contrast Preserving Decolorization method supplied in OpenCV (Lu, Xu, & Jia,2012), and recruited another group of 20 observers to perform the Binary Classification and Material Categorization tasks. The details of the experiment can be found in the Supplementary Material.

## Results

### Experiment 1. Binary Classification

Since “translucent” and “opaque” have been frequently used in the previous studies in the perception of sub-surface scattering materials, we first look at how well observers are able to classify the objects in our image data-set into two categories, “translucent” and “opaque”, and explore how converting images to grayscale affects the binary classification. Our goal is to build a labeled data-set of these photographs for each observer and further examine the relationship between binary classification and semantic attribute ratings (Experiment 2).

#### Procedure

Figure 2 (II) shows the experiment interface. During each trial, the observer was asked to judge whether the material of object shown in the image is either “translucent” or “opaque” through keyboard responses. Prior to the experiment, the observer was introduced to the physical model of light transmission of opaque and translucent materials. Specifically, they were shown an illustration and a brief description of volumetric scattering model and a pair of images rendered with and without sub-surface scattering. To avoid biases, we did not provide photograph examples of translucent materials. 300 images were shown to the observer in a pre-randomized order which was same in both color and grayscale conditions.

#### Results

##### Converting images to grayscale affects the trial-by-trial percent agreement among observers

For analysis, we compute the percent agreement among observers for each trial in the binary classification for both color and grayscale conditions. The percent agreement of an image *i*, *P*_*i*_, is calculated based on following equation (1):

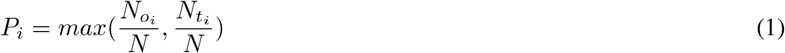

where 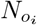 is the number of observers classify the image as “opaque”, 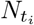 is the number of observers classify the image as “translucent”, and *N* is total number of observers. Figure 3 (I) plots the trial-by-trial percent agreement for both color (red line) and grayscale (blue line) conditions. On the X-axis, the images are ordered from uniformly agreed to be “translucent” (Left) to uniformly agreed to be “opaque” (Right) in the color condition. The figure shows that percent agreement is high for obviously transparent objects ( e.g. glass) and for obviously opaque materials (e.g. metal block), but is low for translucent materials (e.g. scallop). Based on the aggregated data, we can see the ranking of materials based on the percent agreement, from most “translucent” to “opaque”, is largely consistent with the ground-truth. This figure also shows the trial-by-trial percent agreement is different in color and grayscale conditions. For example, some images judged to be “translucent” in color condition are judged to be “opaque” in grayscale.

##### Converting the images to grayscale flips classification labels for some images

We further look at what images are most affected when they are converted to grayscale. Based on the binary classification result, we label an image as “Translucent” (T) if at least 60% of observers classify it as translucent, as “Opaque” (O) if at least 60% of observers classify it as opaque, as “Unsure” (U) if less than 60% of observers classify it as either translucent or opaque. Using these labels, we find that some images flip from “Translucent” in color condition to “Opaque” in grayscale condition, and vice versa. Histogram in Figure 3 (II) shows that 61 images have flipped their classification labels.

**Figure 3:**
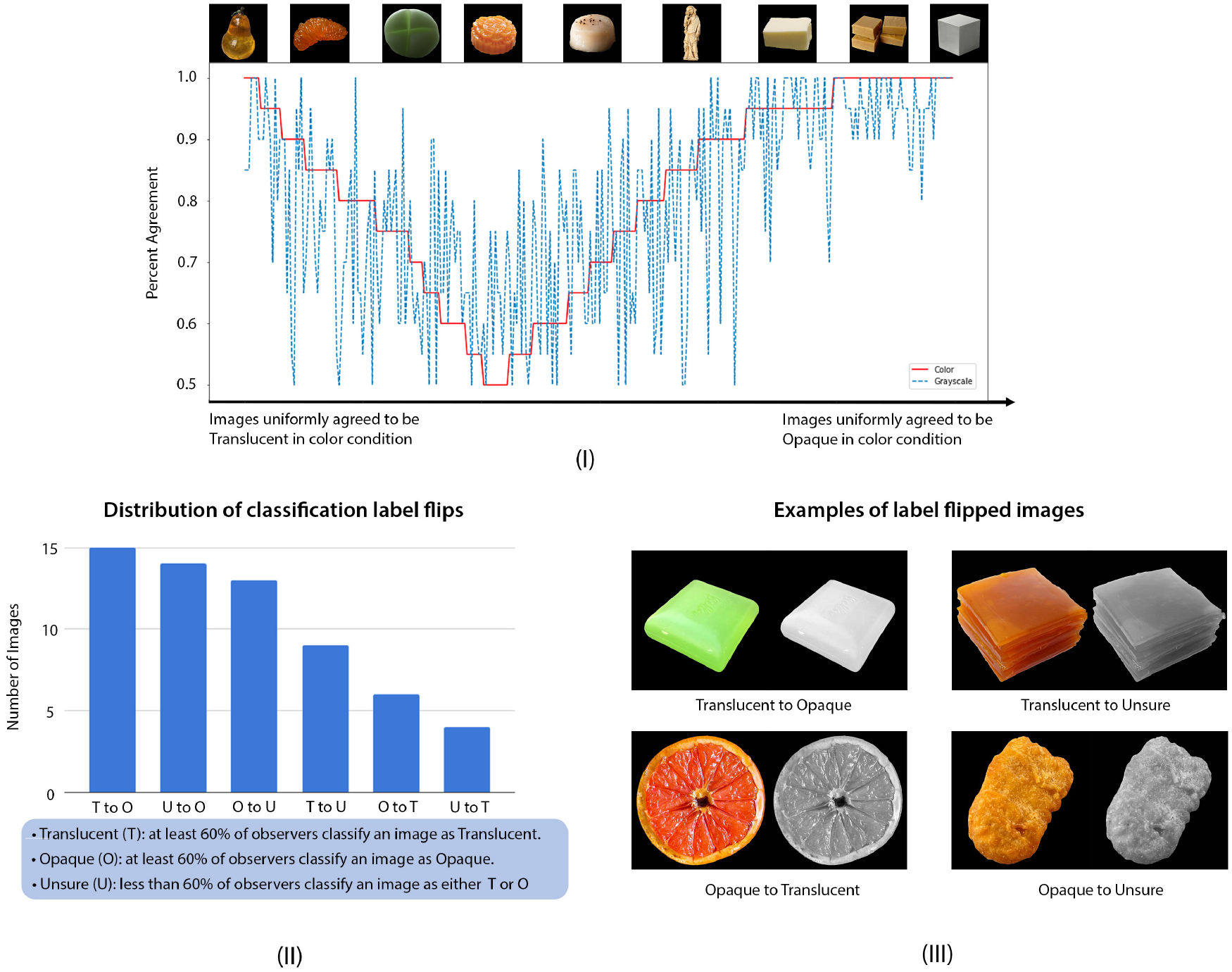
Experiment 1: Effect of color on the percent agreement among observers in binary classification experiment. (I) The trial-by-trial percent agreement of binary translucency classification in color (red) and grayscale (blue) conditions. Top insertions: the example images corresponding to different levels of agreements. (II) Different types of classification label flips when the images are converted to grayscale. The legend at the bottom shows how the labels are defined based on observers’ agreements. For example, “O to U” means the image flips its label from opaque to unsure when it is converted to grayscale. (III) Examples of images that flip the classification label when they are converted to grayscale.

Figure 4 (I) depicts the image-by-image Representational Dissimilarity Matrices (RDMs) (Kriegeskorte, Mur, & Bandettini,2008) based on observers’ judgements in both color and grayscale conditions. Specifically, for each pair of images, we compute normalized Hamming distance based on the 20 observers’ binary classification results. For example, image *i* receives the binary classification label from 20 observers in color condition, and its voting pattern can be represented by an array *a*_*i*_. Similarly, the voting pattern for image *j* in color condition can be represented by *a*_*j*_. Assuming *a*_*i*_ = (1, 0, …, 0), and *a*_*j*_ = (0, 0, …, 1), the normalized Hamming distance between image *i* and *j* is computed by first finding the number of elements at which the arrays *a*_*i*_ and *a*_*j*_ are different, and then dividing it by the number of observers (N = 20). In this example, the normalized Hamming distance is 2 divided by 20, which equals 0.1.

**Figure 4:**
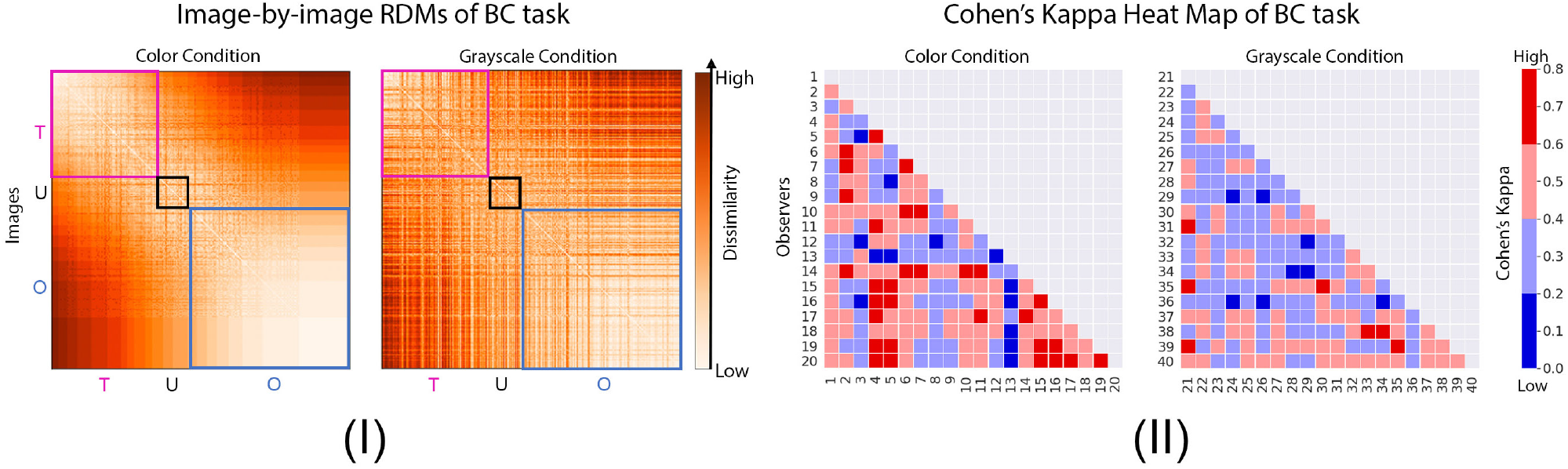
Comparison of individual variability between color and grayscale in binary classification (BC) task. (I) Image-by-image RDMs of the binary classification measured by normalized Hamming distance. The boxed regions represent different categories of images labeled as “Translucent” (T), “Unsure” (U) and “Opaque” (O) in the color condition, which were explained in Figure 3. (II) Person-by-person heat map based on Cohen’s Kappa (*κ*) of the binary classification task (Cohen,1960). Observers in color condition are indexed from 1 to 20, and those in grayscale condition are indexed from 21 to 40.

In Figure 4 (I), the more saturated color in the RDM represents higher dissimilarity (larger normalized Hamming distance). From top to bottom of the axis, we order the images as receiving increasing number of votes of being “opaque” in color condition (same as the X-axis in Figure 3 (I)). The boxed regions represent the images labeled as “Translucent” (T), “Unsure” (U) and “Opaque” (O) in the color condition using the same definition in Figure 3 (II). In color condition, dissimilarity is lower for opaque region (lower right) than the translucent region (upper left). We conduct a one-sided Mann-Whitney U test to compare corresponding regions between color and grayscale and find that all of these regions in grayscale RDM have statistically higher dissimilarity in comparison to those in color RDM (p < 0.0001 for each pair of regions).

##### Converting images to grayscale leads to higher level of disagreements among observers

To investigate individual difference in the results, we compute Cohen’s Kappa (*κ*) as a measurement of inter-observer agreement on the classification of images (Cohen,1960). Cohen’s Kappa can be calculated from the following equation (2):

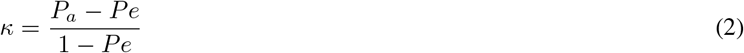

where *P*_*a*_ is the actual observed agreement, and *P*_*e*_ is the chance agreement (McHugh,2012).

In Figure 4 (II), we show Cohen’s Kappa (*κ*) between observers for both conditions. The dark blue and blue cells correspond to low *κ*, meaning that the level of agreement between the pair of observers is none (0 < *κ* <0.20) or minimal (0.21 < *κ* <0.39). The dark red and red cells correspond to relatively higher *κ*, meaning that the level of agreement between the pair of observers is moderate (0.60 < *κ* < 0.79) or weak (0.40 < *κ* < 0.59). From all pairs of observers, we do not observe a strong level of agreement (0.80 < *κ* < 0.90) in either condition. 83.7% and 96.3% of the cells in color and grayscale conditions show minimal and weak level of agreement, suggesting that there is substantial individual difference in the binary classification task. Moreover, there are more pairs of observers reach moderate level of agreement (16.3%) in color condition in comparison to that of the grayscale condition (3.68%). The one-sided Mann Whitney U test comparing the two conditions shows that observers have lower level of agreement when the images are shown in grayscale (p < 0.0001).

#### Discussion

Experiment 1 shows that converting images to grayscale leads to more ambiguity in binary translucency classification for some images, reflected by larger individual variability. Converting to grayscale flips classification labels for some images. We hypothesize that the image cues such as color gradient might be necessary for assessing the translucent appearance, which are absent in grayscale images. Further, color is often associated with object identification, which might also affect the individual’s translucency classification. Without color, object’s identity might be ambiguous and prone to personal interpretation, which might lead to more individual variability in the grayscale condition. For the control experiment with the contrast-preserved version of grayscale images, we obtained the similar results and also observed high ambiguity in the binary translucency classification. The results can be found in the Supplementary Material.

### Experiment 2. Semantic Attribute Rating

In Experiment 2, instead of classifying the material into “translucent” or “opaque”, we ask observers to judge a few commonly used mid-level material attributes associated with translucency with the goal of describing the perceptual space of translucency. Our goal is two-fold: first, we aim to explore to what degree the attribute ratings are affected by converting the images to grayscale; second, we aim to characterize the relationship between the semantic attribute ratings and binary translucency classification, and see whether the selected attributes are related to translucency classification for the individual observers.

#### Procedure

Prior to the main experiment, we conducted a survey after a pilot binary classification experiment (e.g. a shorter experiment done by different observers from the experiment reported here). We asked the observers to describe the image properties or cues they found useful in the binary classification task using adjectives. Based on their responses and previous literature, we selected five perceptual attributes to describe the translucent appearance of materials, including “see-throughness”, “glossiness”, “softness”, “glow”, and “density”. For example, “glow” was first proposed as a cue for translucency by Fleming in (Fleming & Bülthoff,2005). Observers were asked to rate each of these attributes of the material of the object shown in the image on a six-point Likert scale (see Figure 2 (III)). We choose 5 attributes because otherwise the experiment would be very long.

Observers were first asked to carefully read the description of the semantic attributes. On each trial, without time limit, they used a slider to rate each attribute, using 1 for low attribute value and 6 for high attribute value. The order of images was pre-randomized and was different from the binary classification experiment. We define the attributes as follows:

- See-throughness: To what degree the object allows light to penetrate through.
- Glossiness: To what degree the object appears shiny and reflective.
- Softness: To what degree the object is easy to deform.
- Glow: To what degree the object appears to glow light from inside.
- Density: To what degree the object appears to be “dense”.

#### Results

##### Converting to grayscale affects the distribution of attribute ratings for some images

For each image, we plot the distribution of ratings across observers for each attribute. Figure 5 (I) shows some examples of images for which distributions of attribute ratings are affected when they are converted to grayscale, such as the images of a grapefruit and a green soap. To quantify the effect of color on ratings for each image and attribute, we calculate Kullback–Leibler divergence (KL divergence) between the distributions of attribute ratings of color and grayscale conditions. The KL divergence is calculated as following:

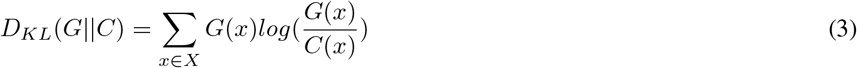

where *G*(*x*) is the probability distribution of ratings for an attribute in grayscale condition, and *C*(*x*) is the distribution of ratings for the attribute in color condition.

**Figure 5:**
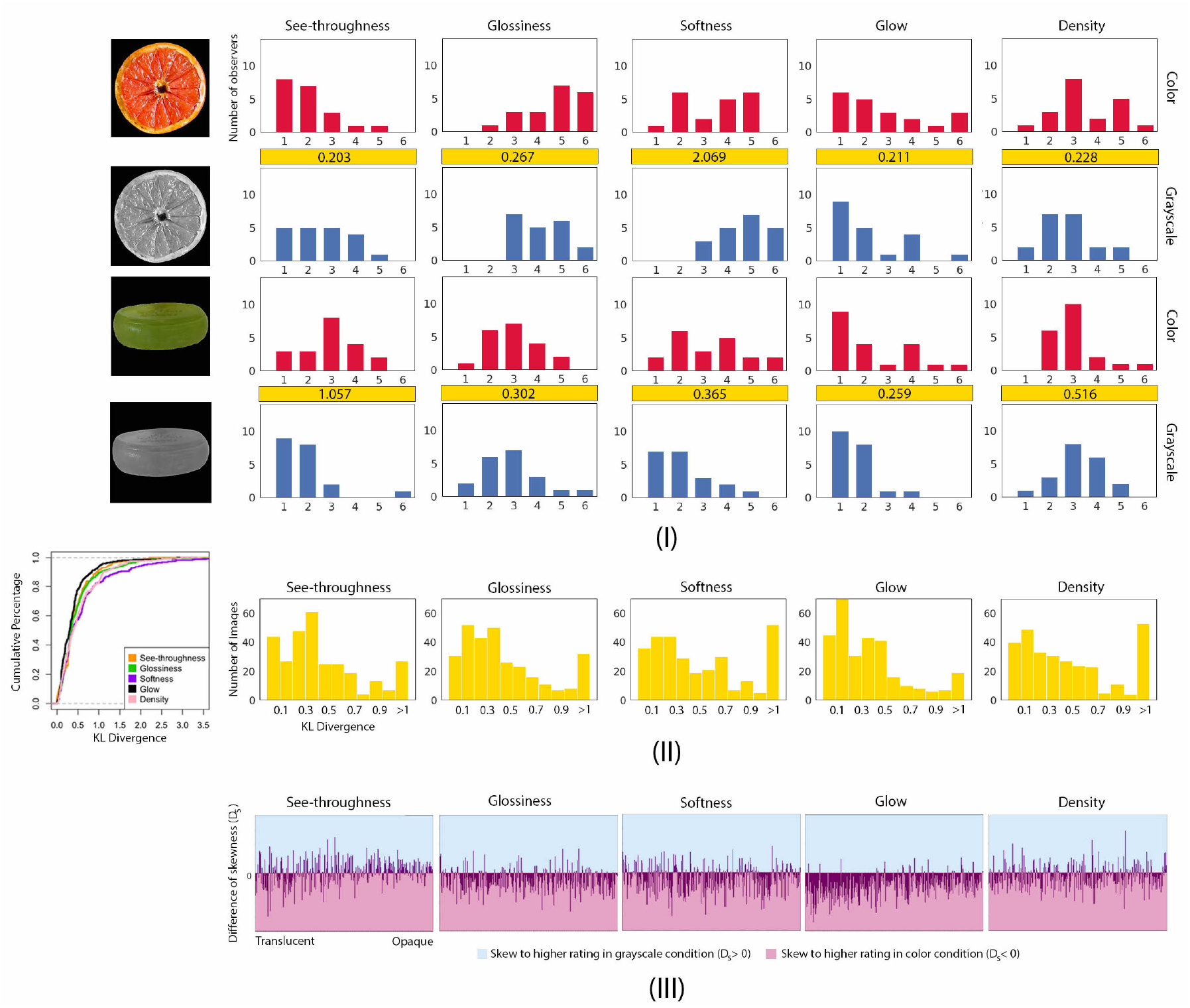
Experiment 2: Effect of converting images to grayscale on material attribute ratings. (I) Examples of the distribution of observers’ ratings of five semantic attributes in color and grayscale conditions. We compute KL divergence between the rating distribution of color condition (top) with that of the grayscale condition (bottom), and show it in the yellow box. (II) Left: The cumulative distribution function of values of KL divergence for each attribute. The X-axis is the value of KL divergence and the Y-axis is the cumulative percentage of images with the corresponding KL divergence value. Right: The distribution of KL divergence of 300 images for each semantic attribute. (III) The effect of converting images to grayscale on the skewness of the distribution of ratings for each attribute. Each line represents the difference between the skewness of the distribution of ratings in color condition and that in grayscale, *D*_*s*_ (see equation 4), for a particular image.

Figure 5 (I) compares the distributions of ratings between color and grayscale images. Figure 5 (II) plots the distribution of KL divergences across all images for each attribute. The figure shows that about 80% of images have relatively small KL divergence and only a small portion of images have high KL divergence. The shape of the distributions of KL divergence is similar across five attributes, with a highly positive skew towards low KL divergence, suggesting that color only weakly affects the attribute ratings for the majority of images. For a small number of images, color has a stronger effect on the ratings, which leads to relatively higher KL divergences. From the two examples, we can see that high value of KL divergence corresponds to visible difference of the shape of the distribution. For the grapefruit images, more observers use relatively high softness ratings in the grayscale condition in comparison to the color condition, resulting in a large KL divergence of 2.069. For the green soap images, observers are more likely to use lower see-throughness ratings in the grayscale condition, also resulting in a moderately high KL divergence of 1.057. Although there is no universal standard for KL divergence, we observe that KL divergence greater than 1 (ie. *D*_*KL*_(*G||C*) > 1) could be an informative threshold indicating significant difference between the distributions of attribute ratings in color and grayscale conditions. Among our stimuli, 9.3%, 12.0%, 19.0%, 7.7% and 17.7% of the images have *D*_*KL*_(*G||C*) > 1 for the distribution of ratings of “see-throughness”, “glossiness”, “softness”, “glow”, and “density”, respectively. Figure 6 shows more examples of images with high (top panel) or low (bottom panel) KL divergence values.

**Figure 6:**
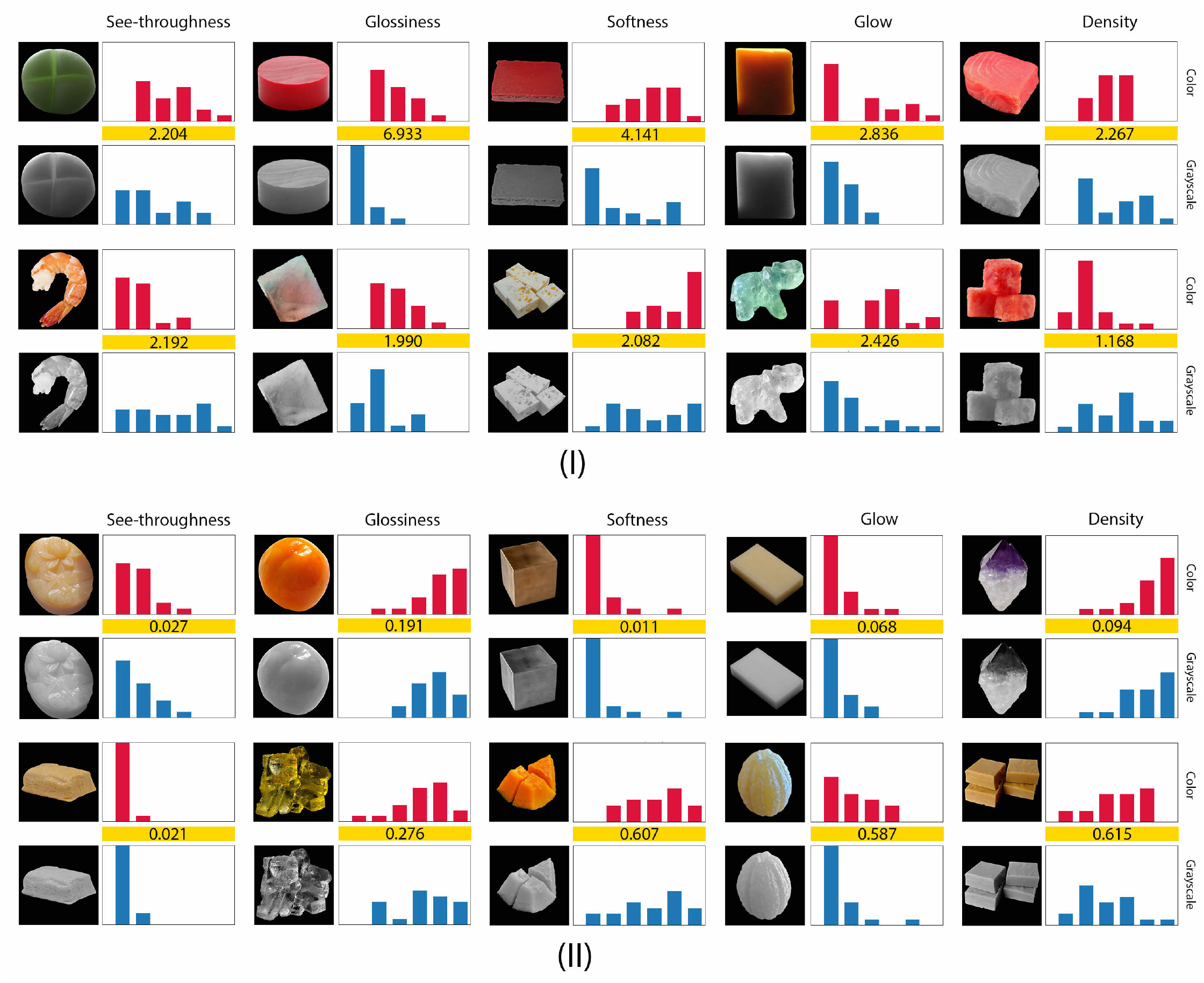
More examples showing the distribution of ratings of each semantic attribute in color and grayscale conditions with their corresponding KL divergence. (I) KL divergences greater than 1. (II) KL divergences less than 1. All axes of the diagrams have the same range as Figure 5.

For a particular image, in order to understand whether the distributions of ratings are shifted towards higher or lower end of the scale when color is removed, we compute the difference in the skewness between color and grayscale, of distribution of attribute ratings, *D*_*s*_, by equation 4.

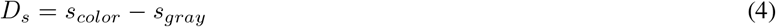

where *s*_*color*_ and *s*_*gray*_ are the skewness of distribution of attribute ratings in color and grayscale condition respectively. Figure 5 (III) shows the *D*_*s*_ of all images in five attribute ratings. The order of X-axis is the same as that in Figure 3 (I). If a line falls in the blue region, it shows that an image has more observers assign higher ratings for that attribute in grayscale condition in comparison to color. We can see that many lines fall into the pink region, meaning that for a great number of images, more observers assign higher ratings to the attributes in color condition. Together, converting images to grayscale causes more observers assign lower ratings of glossiness, softness, glow and density but might cause some opaque objects appear more see-through.

##### The correlations between attribute ratings and perceived level of translucency

Figure 7 plots the average attribute ratings for the images in color and grayscale conditions. On the X-axis of each plot, the images are ordered by the number of votes for being “opaque”, based on the binary classification result in color and grayscale condition respectively. On the left, the images are uniformly agreed to be “translucent”, and on the right, the images are uniformly agreed to be “opaque”. The Y-axis represents the average attribute rating of an image. The figure shows that some attributes’ ratings are significantly correlated with binary classification while others are less correlated. We compute the Kendall’s rank correlation between the mean attribute ratings and the level of translucency based on the binary classification (Experiment 1), and summarize our findings as below:

- Average ratings of see-throughness and glow are significantly correlated with the level of agreement of translucency from the binary classification such that their values decrease as the images are more uniformly agreed to be “opaque”. For see-throughness, the statistical results are *τ* = −0.77, p < 0.05 in color condition, *τ* = −0.74, p < 0.05 in grayscale condition; for glow, we find *τ* = −0.69, p < 0.05 in color condition, and *τ* = −0.6, p < 0.05 for grayscale, suggesting moderate correlation.
- Average rating of glossiness has moderate correlation with binary classification under both conditions (*τ* = −0.44, p < 0.05 in color condition, *τ* = −0.51, p < 0.05 in grayscale condition).
- Average ratings of softness and density are not significantly correlated with the level of agreement of translucency.

**Figure 7:**
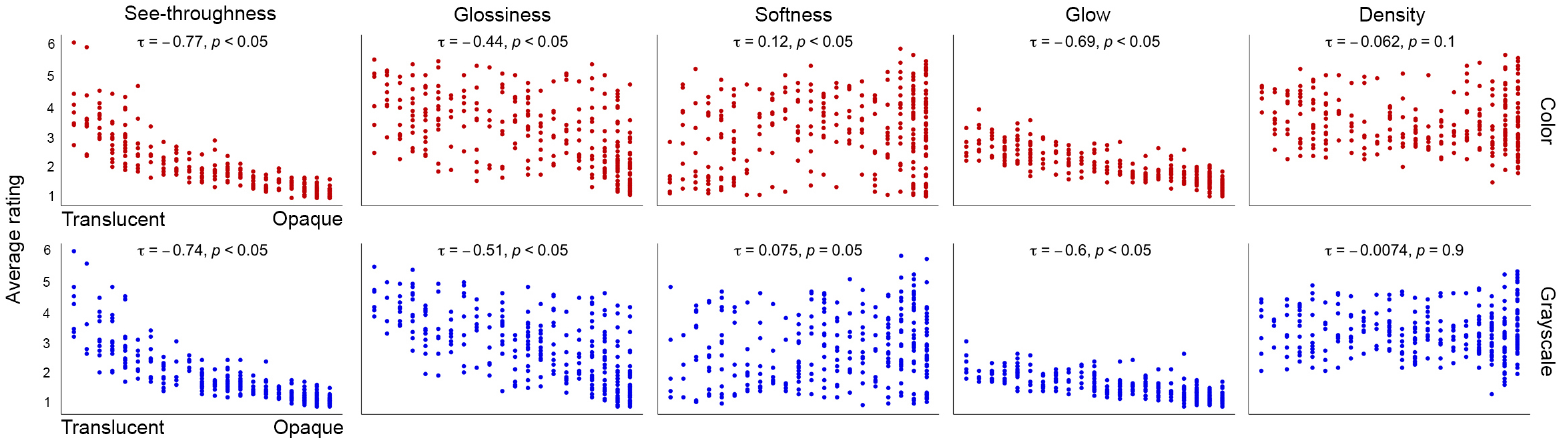
Average see-throughness, glossiness, and glow ratings show correlation with level of translucency resulted from binary classification experiment, whereas average softness and density do not. Each point in the plot represents an image from the data-set. On top of the scatter plots are the Kendall rank correlation coefficient (*τ*) between the average attribute rating and the number of vote for being “opaque”, and the associated p-value at significance level of 95%.

These results suggest see-throughness, glow and glossiness are closely related to how observers classify the objects into “translucent” and “opaque”.

##### Correlation between semantic attributes

We also analyze to what extent the ratings of the attributes are dependent on one another. Figure 8 shows the Kendall rank correlation matrix between the average attribute ratings. Some of the attributes are statistically significantly correlated with Bonferroni correction adjusted alpha level of 0.005 (0.05/10). Overall, the average ratings of see-throughness, glow and glossiness are positively correlated with each other while density and softness are negatively correlated. Specifically, in both color and grayscale conditions, see-throughness is positively correlated with glow (*τ* = 0.67, p < 0.005 in color, and *τ* = 0.61, p < 0.005 in grayscale), and is weakly correlated with glossiness (*τ* = 0.35, p < 0.005 in color, and *τ* = 0.41, p < 0.005 in grayscale); glossiness is also weakly correlated with glow (*τ* = 0.45, p < 0.005 in color, and *τ* = 0.49, p < 0.005 in grayscale); density is highly negatively correlated with softness (*τ* = −0.74, p < 0.005 in color, and *τ* = −0.71, p < 0.005 in grayscale). The correlation between see-throughness and glossiness is consistent with previous findings that glossiness is important in translucency perception and vice versa (Motoyoshi,2010;Gigilashvili, Shi, et al.,2021).

**Figure 8:**
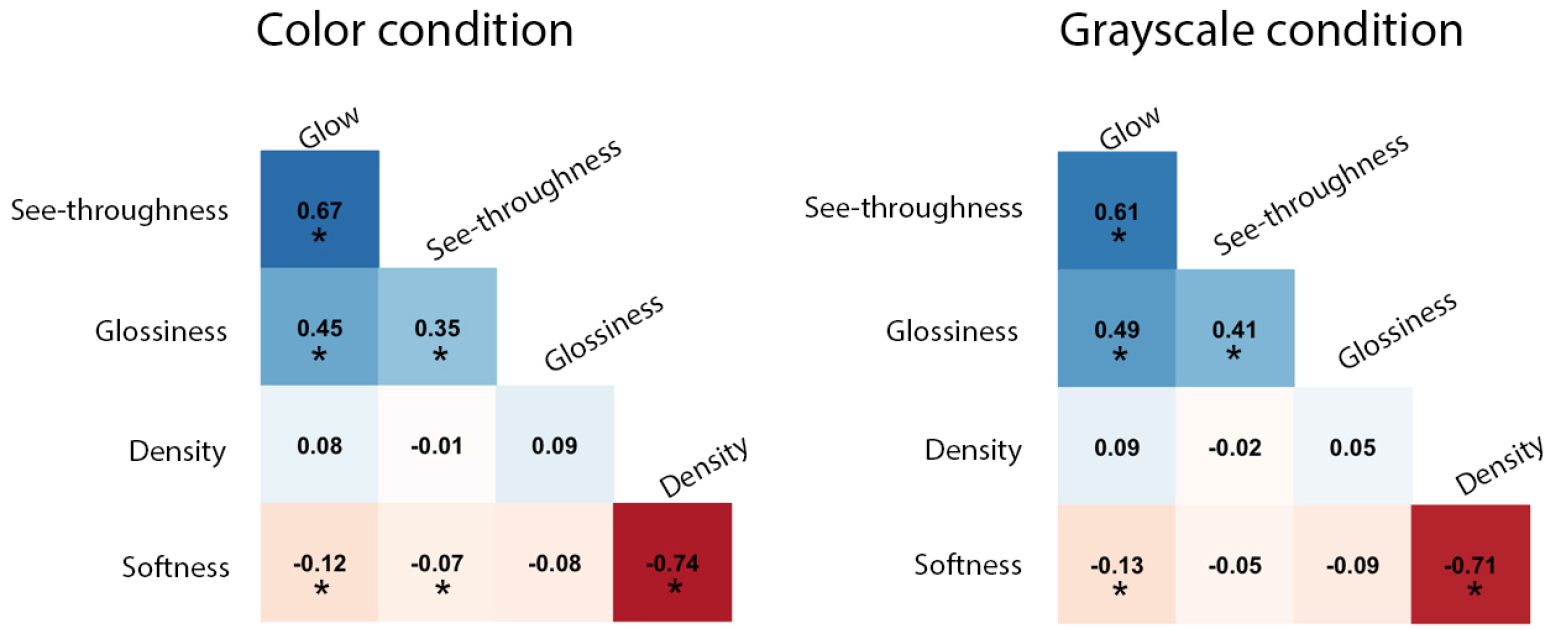
The correlation matrix of average ratings of semantic attributes. Left: Color condition. Right: grayscale condition. The numbers shown in the cells are the Kendall rank correlation coefficients between ratings of two attributes. Blue color represents positive correlation and red color represents negative correlation. * indicates p < 0.005.

##### Clustering of images in the space of perceptual qualities

To understand if the semantic ratings can be used to predict observers’ binary classification, we first compute the mean semantic attribute ratings for each image in both color and grayscale conditions. Then, we use Principal Component Analysis (PCA) to reduce the dimensionality of the rating data. Figure 9 shows the perceptual space of the images with the first two principal components (PCs), where we color the points using the classification labels “Translucent”, “Opaque” and “Unsure” defined in Figure 3 (II). The first two PCs account for approximately 82% of the variance in both color and grayscale conditions. Figure 9 (I) shows that the images can be clustered into “Translucent” and “Opaque” in the 2D perceptual space. In the right panels of Figure 9 (I), the arrow-ed vectors describe how much each semantic attribute contributes to a particular PC. Large loading (either positive or negative) indicates the attribute having a strong relationship with a particular PC, and the sign of loading indicates whether an attribute and a PC are positively or negatively correlated. In the color condition, see-throughness, glow, and glossiness have slightly higher weights in first PC (PC1), and softness and density have large loadings on second PC (PC2). In the grayscale condition, see-throughness, glow, and glossiness primarily contribute to PC1, while softness and density primarily contribute to PC2. In both conditions, we see that the loading vectors of see-throughness, glow, and glossiness are orthogonal to the direction spanned by density and softness, suggesting that the light transmission related attributes are highly correlated with each other and softness and density are anti-correlated, confirming the results shown in Figure 8. Figure 9 shows that the direction spanned by see-throughness, glow, and glossiness separates “Translucent” cluster from the “Opaque”, suggesting that these three attributes could be indicative of “Translucent” materials labeled in the binary classification task.

**Figure 9:**
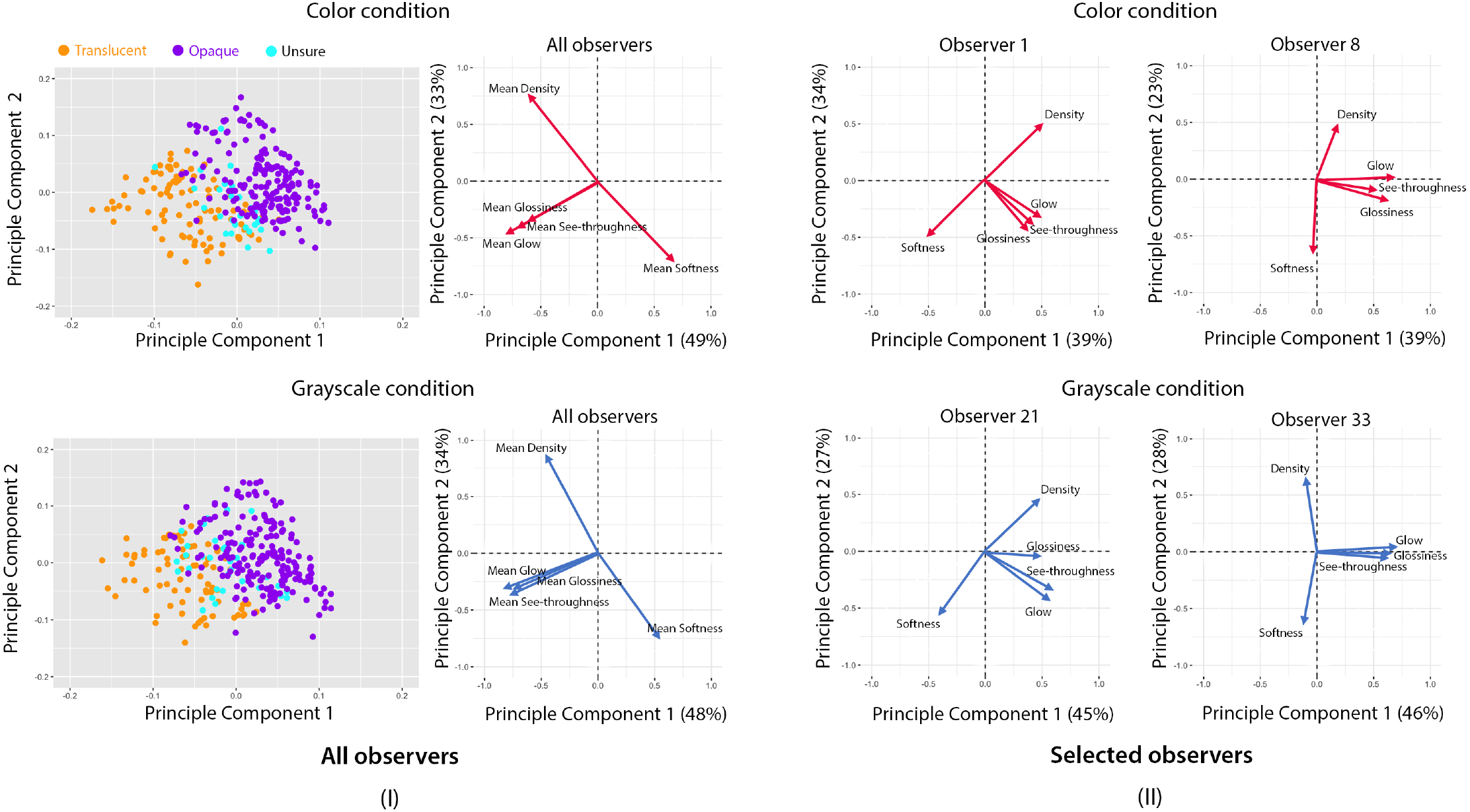
(I) A principal component analysis (PCA) of the mean attribute ratings in color and grayscale conditions. Left: PCA scatter-plots of 300 images in color and grayscale conditions. Right: The loadings of five semantic attributes. The axis represents the loading on the first or second principal component, with the explained variance shown in parentheses. (II) The loading of attribute of selected observers in color and grayscale conditions.

To look at the results on the individual level, we use each observer’s attribute ratings to perform PCA analysis based on polychoric correlation (Revelle,2020). Common across all observers, similar as results from mean ratings, we find that the loadings of density and softness are mostly orthogonal from those spanned by see-throughness, glow and glossiness. Figure 9 (II) shows the loading of each attribute on the first two PCs and the correlation among them vary somewhat across observers.

##### See-throughness, glossiness and glow can be used to predict observers’ binary translucency classification

While considering the presence of individual variability in semantic attribute rating and binary classification tasks, we aim to understand the relationship between them. For each observer, we use their five attribute ratings as features, and perform a logistic regression with 3-fold cross validation to predict the observer’s binary classification of an image. Figure 10 (I) shows that using logistic regression achieves 80% prediction accuracy for most observers in both color and grayscale conditions. While running the logistic regression using three different train-test splits for each observer (3-fold cross-validation), we record the attributes that are statistically significant in predicting the binary classification at significant level of 95%. For each train-test split, we count the number of observers who reach statistical significance for each attribute, and repeat this three times. Thus, for each attribute, we obtain 3 measurements representing the number of observers reach statistical significance. Figure 10 (II) plots the median value of the measurements for each attribute. In both conditions, see-throughness is a statistically significant predictor for all observers, and glow and glossiness are significant features for at least 30% of observers. Softness and density are statistically insignificant predictors for most observers. Although softness and density themselves may not be strong predictors for the binary translucency classification, they might be used together with other attributes in binary classification for

**Figure 10:**
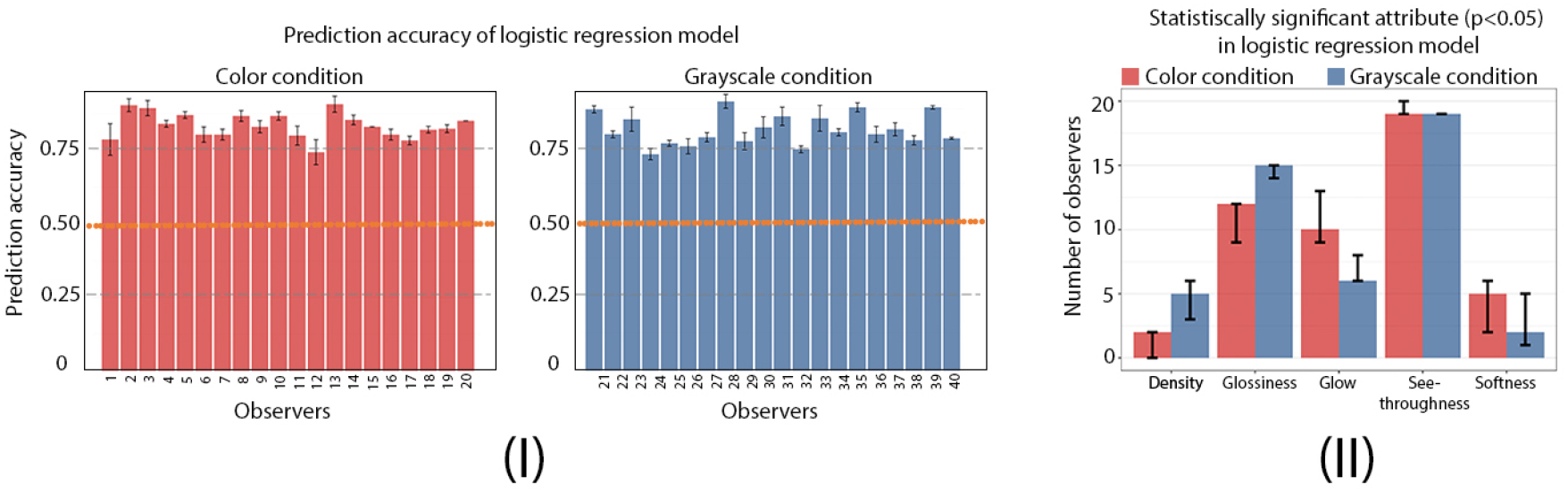
Prediction of individual observers’ binary classification from semantic attribute ratings using the logistic regression. (I) Prediction accuracy for observers in color and grayscale conditions. The error bar indicates the standard deviation of the prediction accuracy of the 3-fold cross-validation. Orange line corresponds to prediction by chance (50%). (II) The median number of observers in the 3 cross-validations that reaches statistical significance (p < 0.05) in the logistic regression for each semantic attribute. The error bar shows the difference between the maximum and minimum number of observers in the 3 cross-validations some observers.

#### Discussion

In Experiment 2, we find color differentially affects ratings of certain images and attributes. It is possible that these effects depend on specific lighting condition and the object’s intrinsic material properties. Figure 6 provides more example images, demonstrating high and low KL divergences, and shows how converting images to grayscale affects the distributions of semantic attribute ratings. We conjecture that low KL divergence between color and grayscale ratings of a particular attribute indicates the easiness of perceiving this attribute and recognizing the material category from the image (see Figure 6 (II)).

Based on both group and individual results, we see that see-throughness, glossiness and glow are correlated, and that they are statistically significant features to predict the observers’ binary translucency classification in both color and grayscale conditions. For some observers, see-throughness alone is an effective feature to predict whether an image is judged as “translucent”. For others, the combination of see-throughness with other semantic attributes is required. Even though density and softness might not contribute to the perception of translucency, we find removing color affects their distributions of ratings. As shown in Figure 5, there are more images having high KL divergence in the ratings of density and softness, suggesting that for some images, converting them to grayscale effectively influences observers’ estimation of these two attributes. Finally, our list of attributes might not be complete. But we do not think adding additional attributes will change the results such that see-throughness, glow, and glossiness are important in the perception of translucency. In the next experiment, we are interested in exploring the likelihood that observers’ attribute ratings are related to material recognition.

### Experiment 3.Material Categorization

In this experiment, we measure material categorization beyond binary classification using the same image data-set, investigate whether and how converting images to grayscale affects categorization, and explore the relationship between semantic attribute ratings and material classes.

#### Procedure

To decide on which categories to include in the options provided to the observers, we first asked the authors in this paper to independently write down the material names for each image in our data-set and generated a list of categories based on the consensus. Figure 2 (IV) shows the experiment interface. On each trial, without any time limit, observers used the radio button to select the category of the material that is most appropriate to the object shown in the image from 18 options, including Food in general, Food/cheese, Food/fruit/vegetables, Food/gummi/jelly, Food/meat/seafood, Food/candy/candy, Food/shaved ice/ice cream/cream, Crystal/quartz/mineral/jade, Glass, Ivory, Marble/stone/concrete, Plastic/synthetic, Rubber, Soap, Wax, Wood, Chalk and Metal. Excluding the images that contain object with uncommon materials, we selected 287 images from the data set. The order of images was pre-randomized and was different from the binary classification and semantic attribute rating experiments. For both color and grayscale conditions, we asked 15 observers who participated the binary classification and semantic attribute rating experiments to perform this task.

#### Results

##### Effect of color on material categorization

Figure 11 shows the number of observers misjudge the material category of an image in color and grayscale conditions. Each radar chart contains the images belonging to a specific ground-truth material category, and the distance from the data point to the center represents the number of observers select the incorrect material for an image. Instead of showing the radar chart for all 18 categories in the experiment, we discard “chalk” and “wood”, which only have few images in the data-set, and combine the rest to the following 8 major categories: “Food in general”, “Soap”, “Marble/stone/concrete/ivory”, “Glass”, “Crystal/quartz/mineral/jade”, “Plastic/synthetic/rubber”, “Wax” and “Metal”. Overall, observers are more likely to misjudge material categories of objects in grayscale images in comparison to color images, which is confirmed by our statistic test (one-sided Mann-Whitney U test, p < 0.005). The error rates are especially high for the materials such as “Food in general”, “Soap”, and “Wax”.

**Figure 11:**
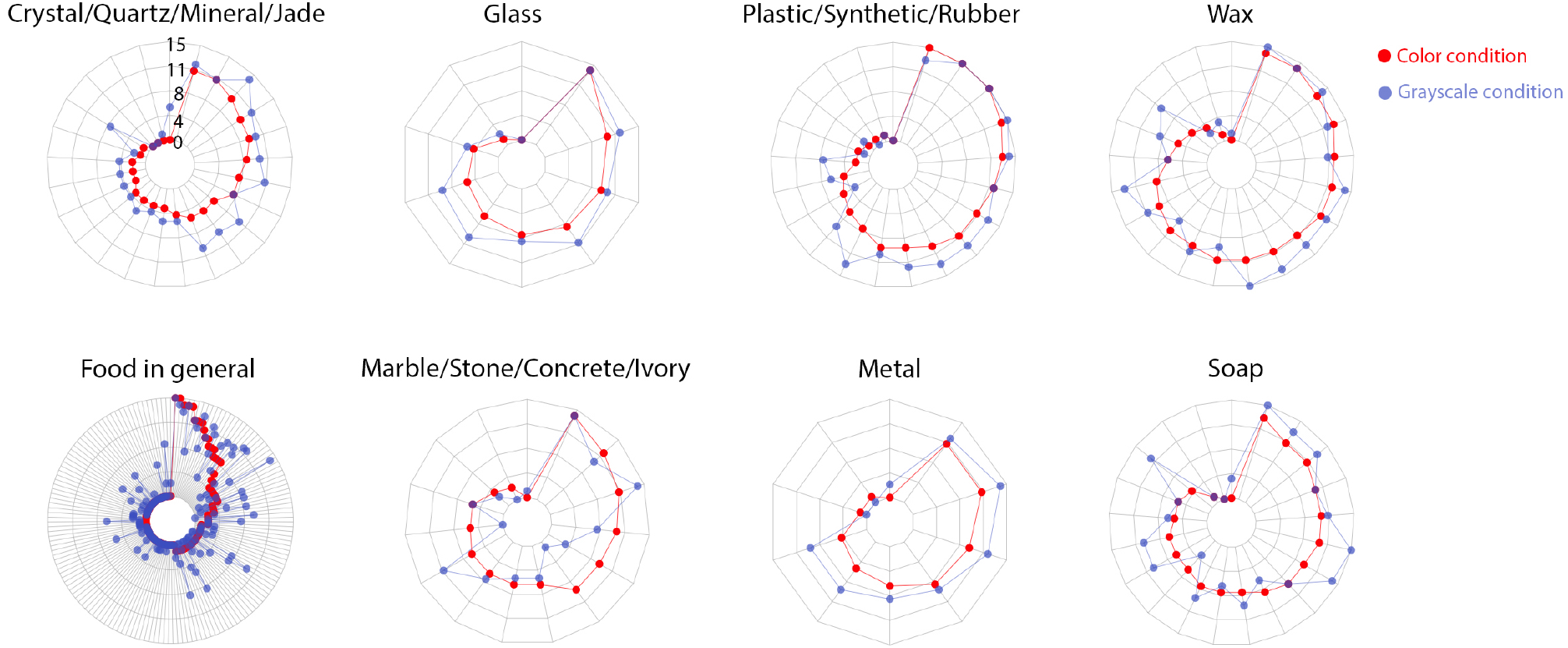
Experiment 3: Radar charts showing the number of observers misjudge the material category of the image in color and grayscale conditions. Each radar chart represents the images belonging to the same ground-truth material category. The distance from the point to the center depicts the number of observers who misjudge the material category of a corresponding image in color (red) or grayscale (blue) condition. All charts use the same range.

In the absence of color, observers were especially poor at identifying “Food”. After we obtain the material classifications from observers, we manually regroup the material classes, “Soap”, “Marble/stone/concrete/ivory”, “Glass”, “Crystal/quartz/mineral/jade”, “Plastic/synthetic/rubber”, “Wax” and “Metal”, as “Non-food” and the rest as ‘Food in general’, and examine observers’ material classification under this binary setting. Figure 12 (I) shows the trial-by-trial material categorization in both conditions. The X-axis is ordered by the images uniformly classified as “Food in general” (Left) to uniformly classified as “Non-food” (Right) in the color condition, and the stacked bar shows the number of observers judge the material as food or non-food. The figure shows that trial-by-trial classifications of food versus non-food are different for some images in color and grayscale. Some objects uniformly agreed as “Food in general” in the color condition are judged by more observers to be “Non-food” in grayscale. To relate to our previous experiments, we notice that some of these misjudged images tend to be translucent. For example, some observers reckoned the watermelon chunk as “Crystal/quartz/mineral/jade”, in grayscale condition. On the other hand, the transparent plastic green lemon is judged by most observer as “Non-food” in color condition but judged as “Food” by some observers in grayscale. Figure 12 (II) shows more examples of images and their distribution of judged categories. Even with color images, observers do not always agree on the material categories, and there is more disagreement in grayscale. For the unfamiliar materials, such as the resin cube in Figure 12 (II), which can take any shape and color, observers are more likely to misjudge the material category in both color and grayscale conditions. This might be caused by their individual difference in object recognition and the association between color and object identity.

**Figure 12:**
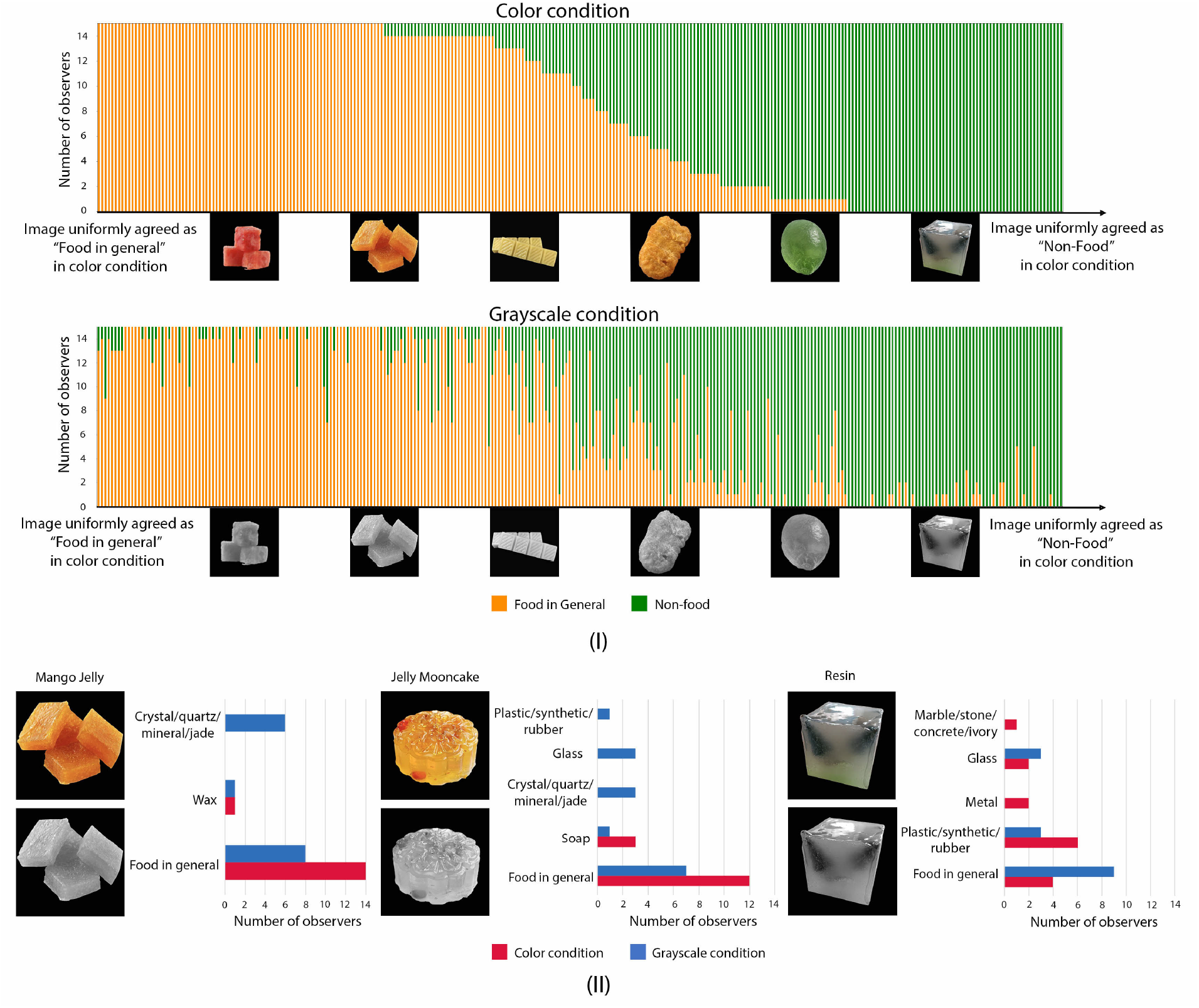
Experiment 3: Observers do not reach uniform agreement on food and non-food categorization of some materials. Some images uniformly judged as “Food” in color condition, are judged as “Non-food” in grayscale condition, and vice versa. (I) Trial-by-trial categorization of image as “Food in general” or “Non-food”. Material categories, including “Crystal/quartz/mineral/jade”, “Marble/stone/concrete/ivory”, “Glass”, “Plastic/synthetic/rubber”, “Soap”, “Wax” and “Metal”, are further generalized as “Non-food”. The rest are generalized as “Food in general”. (II) Examples of images with different material categorization results from observers in color and grayscale conditions. The histograms show the distributions of material category judged by the observers. For example, mango jelly was uniformly judged as “Food in general” in color condition (red bar), but was judged as “Crystal/quartz/mineral/jade” by some observers in grayscale condition (blue bar).

##### Effect of color on observers’ disagreements in material categorization

On the level of individuals, observers are more likely to judge differently in the grayscale condition. Figure 13 plots the person-by-person RDMs, which use normalized Hamming Distance to measure to what degree a pair of observers are different in material categorization. We find that the two RDMs are significantly different.

**Figure 13:**
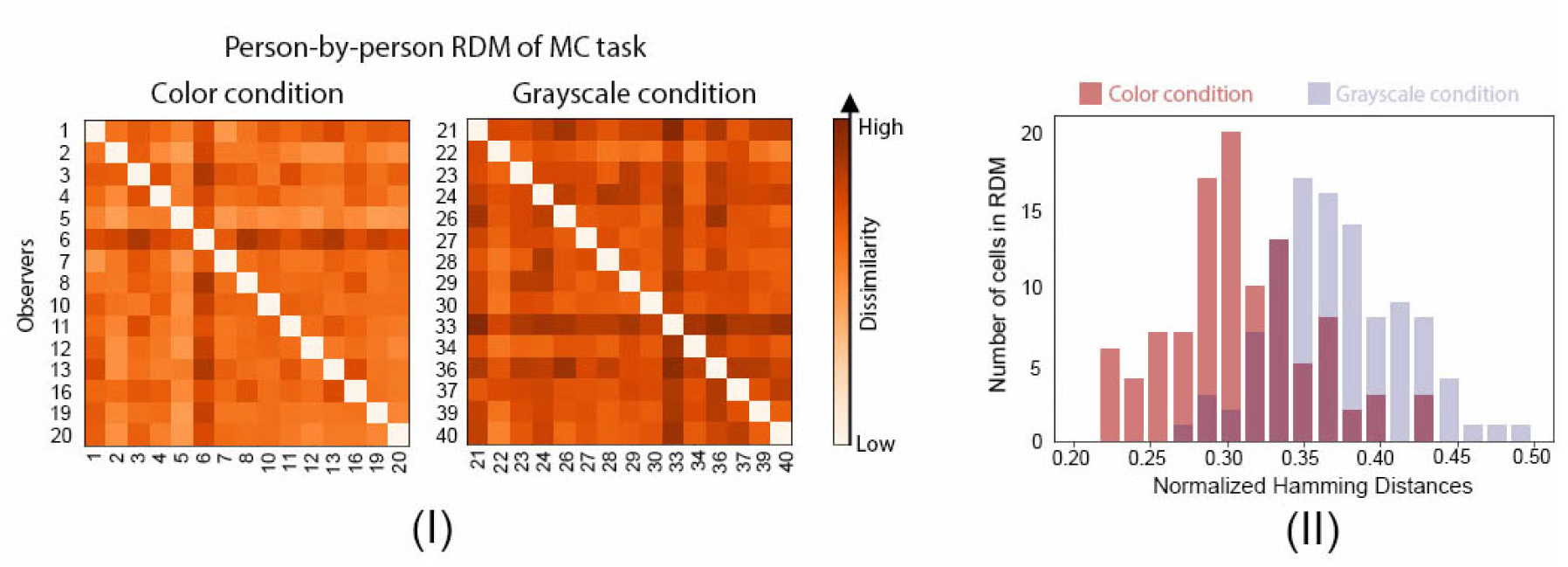
Experiment 3: Converting images to grayscale results in more observer disagreements in material categorization (MC) task. (I) Person-by-person RDMs of Experiment 3 in color and grayscale conditions. For each condition, 15 observers who completed the binary classification and semantic attribute rating experiments performed this task. The axes represent individual observers with the same observer index shown in Figure 4 (II). The dissimilarity is computed from observers’ material classification results of 287 images using normalized Hamming distance (see Experiment 1 for how this is computed). More saturated-color cells correspond to higher dissimilarity between two observers, indicating that they are more likely to disagree. (II) The comparison of distributions of the normalized Hamming distances in color and grayscale conditions.

Specifically, the grayscale condition RDM is more likely to have high dissimilarity values in comparison to the color condition RDM (one-sided Mann-Whitney U test, p < 0.0001).

##### Relationship between material categorization and semantic attribute ratings for individual observers

Figure 14 shows the t-SNE (Maaten & Hinton,2008) plots of images using five semantic attribute ratings as features, colored with the observer’s choice of the material category, for selected observers. We can see that the images that are closer in the embedding spanned by the semantic attribute ratings are likely to be classified as similar materials. It is possible that observers involuntarily use material recognition as an intermediate step to estimate the material properties. Also, we notice that material categories, such as “Crystal/quartz/mineral/jade”, “Marble/stone/concrete/ivory”, and “Metal”, are close to each other in the t-SNE embedding. Observers might not be able to meticulously assign distinctive attribute ratings to them. Further, our results show that it is possible to separate the materials into “Food” and “Non-food” from the t-SNE plot derived from semantic attribute ratings. It is interesting to see the perceptual space extracted from semantic ratings not only can cluster the images into binary classes “translucency” and “opaque”, but also can cluster them into “Food” and “Non-food”, suggesting that there might be common representations between food-related material classification and translucency perception.

**Figure 14:**
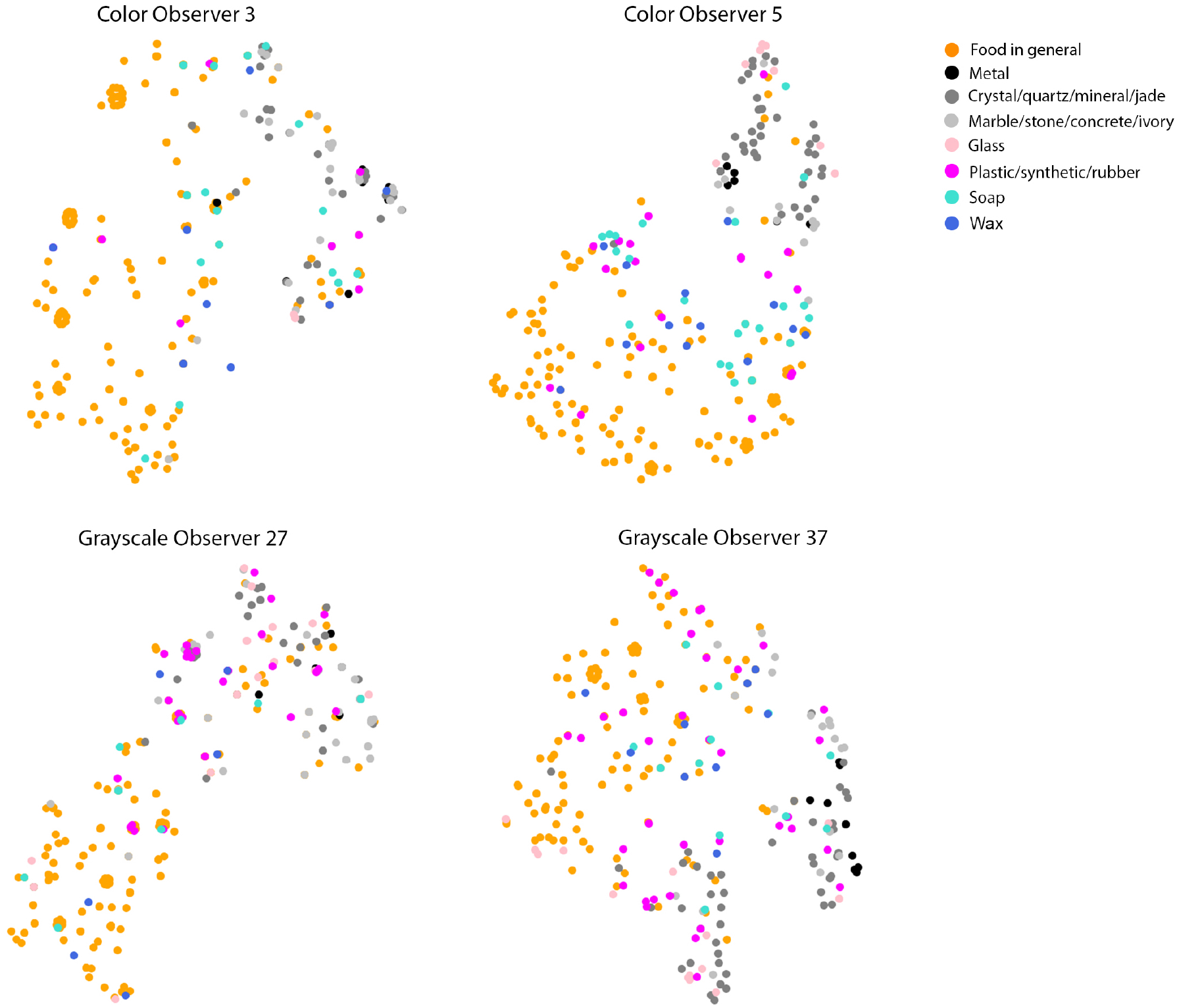
t-SNE (Maaten & Hinton,2008) embeddings (perplexity = 15) of individual observer’s material attribute ratings show clusters emerge corresponding to the perceived material categories. It is easy to see “Food” is clustered away from “Non-food” for most observers. Hard materials such as glass and crystals are also clustered together. Top panels are two observers’ embeddings in color condition and the bottom panels are two observers’ embeddings in grayscale.

#### Discussion

When color is removed, we notice that observers are incapable of recognizing the ground-truth material category of some objects. Particularly, observers often misjudge “Food” as “Non-food” for the physically translucent objects in grayscale, suggesting that color could be a cue that is associated with food recognition. We also obtained similar results for the control experiment with the contrast-preserved version of grayscale images. The results can be found in the Supplementary Material. To relate the results in material categorization to our previous experiments, we find that the images judged to be the same material class by the observer are close to each other in the individual t-SNE embedding of ratings, providing the hint that observers might rely on material recognition to estimate the semantic attributes related to translucency. However, we need future work to investigate the causal directionality of the relationship between semantic attributes and recognition of material categories. It also remains to be investigated how the perception of food is related to the perception of translucency.

## General Discussion

Our study is the first to measure the effect of color on translucency perception using photographs of real-world objects. We find converting images to grayscale affects both material property estimation and recognition of translucent objects. In Experiment 1, we find that converting images to grayscale affects the trial-by-trial percent agreement among observers of whether an object is “translucent” or “opaque”. We also find that there are more disagreements among observers in the grayscale condition. Also, converting images to grayscale causes some objects in the images to change their class label from “Translucent” to “Opaque” and vice versa. In Experiment 2, we find that converting images to grayscale substantially affects the distributions of semantic attribute ratings for some image/attribute combinations. In particular, removing color causes the distributions of ratings of glossiness, softness, glow and density to skew towards lower values. We also find the ratings of see-throughness, glossiness, and glow are moderately correlated with the degree of translucency derived from binary classification. We further show that the ratings of the attributes can be used to predict individual observers’ binary classification, indicating a combination of these attributes might be used by the observers to perceive translucency. However, it is still insufficient to conclude the causal relationship between the semantic attribute ratings and the binary classification. In addition, common across all observers, we discover that see-throughness, glow, and glossiness are positively correlated, and density and softness are negatively correlated. Finally, in Experiment 3, we find observers are more likely to misjudge the material categories in comparison to ground-truth for grayscale images. This effect is especially strong when observers are judging the images of food versus non-food categories.

Overall, we find that removing color has a differential effect on images in our data-set, meaning that some images and material attributes are more affected than others. In the mean time, we find material categorization is related to semantic attribute ratings, which is consistent with previous studies (Fleming et al.,2013;Zuijlen et al.,2020). However, different from previous work, we analyze the data from the level of individual observers, and discover that converting images to grayscale results in more inter-observer disagreements.

### Effect of color: high-level association versus low-level image cues

We conjecture that color affects translucency perception in two possible pathways. One is through high-level association between color, object identity, and individual understanding of translucency. Specifically, color might affect recognizing an object through association (e.g. a red shrimp is usually cooked). In turn, there might be an association between object identity and whether an observer believes it is translucent (e.g. a cooked shrimp is opaque). Therefore, when color is removed, it is likely for some observers to judge a grayscale shrimp to be raw and therefore, translucent.

Observer’s misjudgement of “Food” versus “Non-food” in Experiment 3 might be due to the association between color and object’s category. When color is removed, observers might rely more on other low-level cues and object’s shape to judge its category. How an observer associates color with material category might depend on their experiences and cultural backgrounds as well. Previous work has hinted color affects object recognition (Tanaka, Weiskopf, & Williams,2001). However, the complex relation among color, shape, object identity, and material categorization remains to be investigated.

Another pathway is through low-level image cues. Previous work suggested color-related image statistics such as saturation might affect the perception of translucency and wetness (Fleming & Bülthoff,2005;Sawayama et al.,2017). Inspired by earlier observations (Xiao et al.,2012;Nakano et al.,2009), we notice that manipulating the Pearson correlation between the luminance and saturation affects the translucent appearance of some materials. In the case of the wax cube image shown in Figure 15, the center of the cube looks deeply yellow because light passes through a greater distance, but it looks pale yellow when light passes through the thin edges. This attenuation effect is manifested in the saturation dimension. For translucent materials, this variation of saturation at thin and thick parts of an object can be an important cue for the translucent appearance. The spatial variation of saturation can be negatively correlated with that of luminance in translucent object. According to the optical extinction process in a translucent medium, a decrease in radiance due to a large amount of light extinction should be accompanied by an increase in color saturation. Previous work has shown that manipulating the correlation statistics between color and lightness can impact image translucency transfer (Todo, Yatagawa, Sawayama, Dobashi, & Kakimoto,2019).

**Figure 15:**
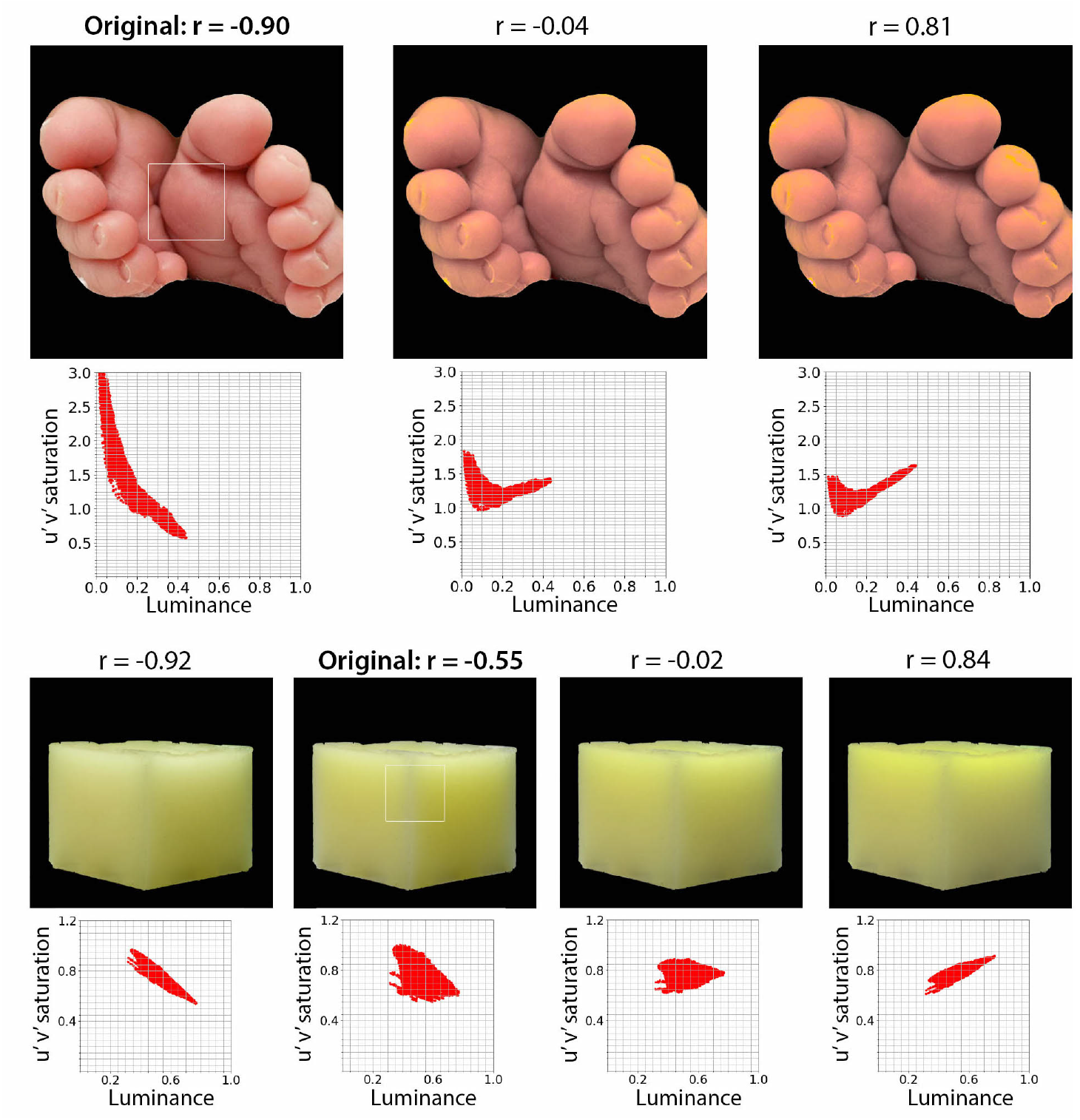
Correlation between saturation and luminance. The images of baby’s feet and wax (“Original” in the figure) were transformed by manipulating the correlation between CIE u’v’ saturation and luminance. The scatter plot under each image indicates pixel values of u’v’ saturation and luminance for the region pointed by the white square on the original image. The Pearson correlation coefficient (r) of the distribution is shown on the top of each image. The transformation has been applied only to the saturation distribution, not the luminance distribution. Specifically, the pixel saturation has been modulated by linearly blending normalized distributions of saturation and luminance. When the weighting of luminance distribution increases, the output saturation changes to the direction of positive correlation. The mean and standard deviation of the original saturation distribution are kept constant.

Figure 15 shows the effect of manipulating the correlation between CIE u’v’ saturation and luminance on the appearance of images. The correlations for two original translucent images (“baby’s feet” and “microcrystalline wax”) show negative correlations (“Original” in Figure 15). When we manipulate the saturation distribution to decrease the negative correlation between saturation and luminance, the translucent appearance on the “baby’s feet” image drastically reduced (center and right in Figure 15), consistent with the notion of previous literature (Fleming & Bülthoff,2005;Xiao et al.,2012). In contrast, when we manipulate the correlation for the “microcrystalline wax” image, both the negative and positive correlations show translucent appearance. Specifically, the negative correlation makes the edge of the wax cube appear diluted in color, therefore giving an impression of “icy” translucent. On the other hand, the positive correlation makes the center of the cube more yellowish and gives an impression of “glow”. Therefore, the image with the positive correlation appears “glow” or “warm” translucent. This finding is also mentioned in earlier translucency work by Fleming et al. (Fleming & Bülthoff,2005), indicating the positive correlation between the saturation and luminance is also plausible. The correlation between saturation and luminance might play a role in translucency perception, but the way is complex.

We also notice that the Chroma channel provide important cues for translucent appearance. Figure 16 shows some examples of label flipped images from the binary translucency classification task (see Figure 3 (II)) in CIELCh color space. For some images whose label flipped from translucent to opaque, such as the pink wax, the gradient in the Chroma channel varies significantly from the center to the sharp edges, creating an impression of glow from the inner area of the object. This gradient is less obvious in the Lightness channel. On the other hand, for some images whose label flipped from opaque to translucent, such as the grapefruit, the image of the Chroma channel appears to capture the light reflection layer of the image and by itself looks more opaque than the Lightness channel.

**Figure 16:**
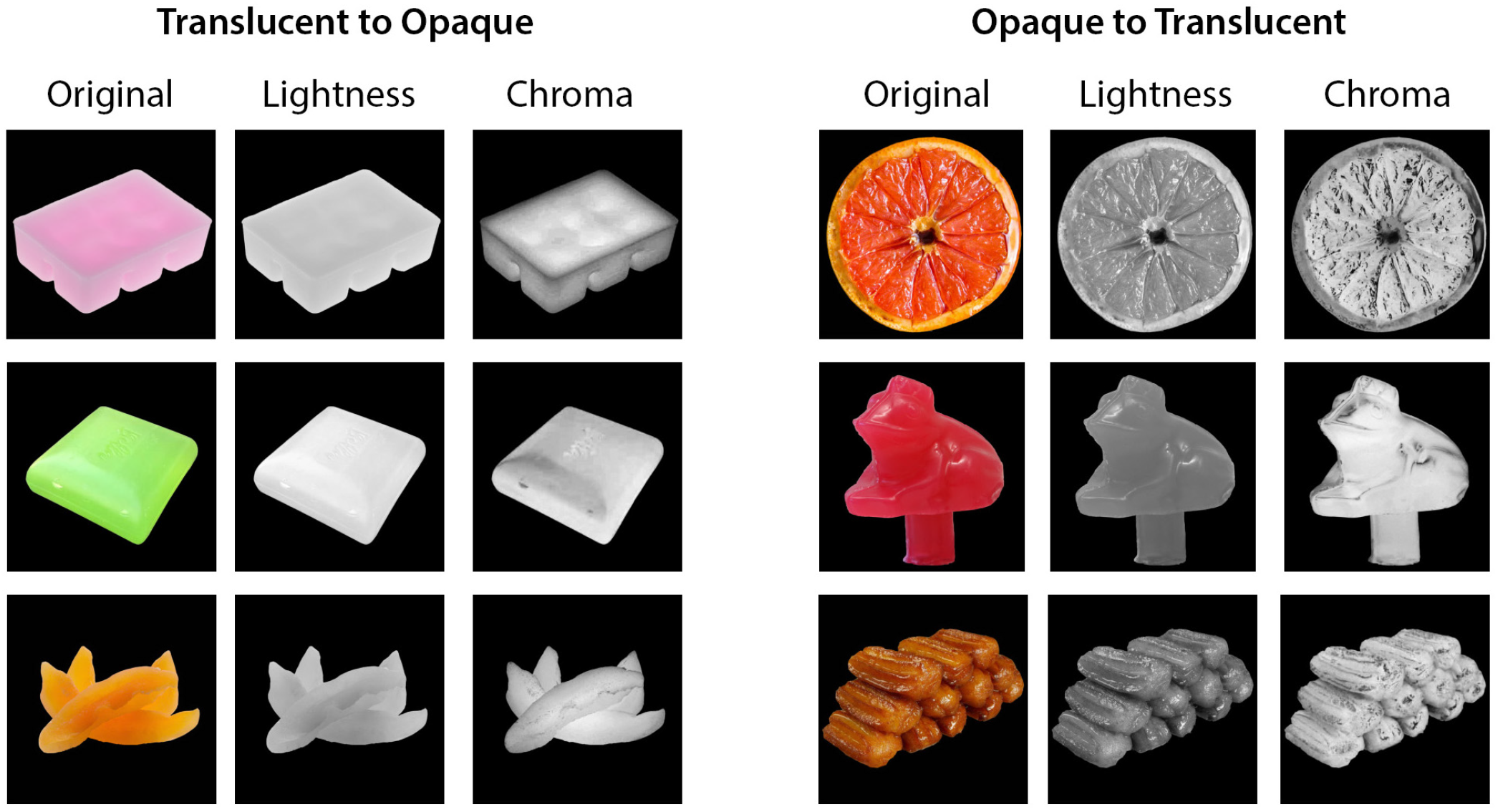
The Lightness and Chroma channels of the “label flipped” images selected from Figure 3 (II). Left: examples of “Translucent to Opaque”. Right: examples of “Opaque to Translucent”. For each example, the orginal RGB images are converted to the CIELCh color space, and the Lightness and Chroma channels are extracted.

### Individual differences

We discover that there are substantial individual differences in our tasks, which we believe reflect the underlying uncertainty of the observers’ decisions in the experiments. We find converting images to grayscale results in more individual variability in both Experiment 1 (Figure 4(II)) and Experiment 3 (Figure 13). Previous works in material perception has not systematically investigated individual difference as those in color vision (Brainard & Hurlbert,2015;Mollon, Bosten, Peterzell, & Webster,2017) except a recent work on translucency perception (Gigilashvili et al.,2020), which discusses the challenges of ambiguity in the concept of translucency perception. In our experiments, individual difference can potentially emerge at various processing stages. We summarize as the following:

- Individual difference in the high-level understanding of “translucency”. We discover that there is individual difference in binary classification experiment, especially when the materials are physically translucent. It is possible the inter-observer differences we observed in the binary classification might be attributed to the subjective understanding of what translucency means. Some observers might not be fully aware of the physical process of sub-surface scattering and tend to label most objects as “opaque”, while others might use their knowledge of the physical process of scattering in the judgements. This might result in semantic ambiguity in the binary translucency classification. We mitigate this by seeking a perceptual space using translucency related semantic attributes. We find that even though there are still individual variability in the ratings, individual observers’ own ratings are correlated with their binary classification results. This shows that although observers might have different semantic-level understanding of “translucency”, there is commonality in how their material attribute rating, a mid-level representation, is related to their binary translucency classification.
- Individual difference in prior experience of interacting with materials. We find individual difference in material categorization even in the color condition, especially in judging “Food” versus “Non-food” of translucent objects (see Figure 12). One reason might be that observers have different experiences and knowledge of certain materials (e.g. Asian observers might be more familiar with mochi and mooncake). Future work is needed to understand how life experience and culture background affect recognition of material categories and explore the plausibility of their relationship to translucency perception.
- Individual difference in low-level perception. Difference in the low-level perception, such as which image regions observers attend to and what local image cues observers use, might affect observers’ judgements (Nagai et al.,2013). This type of individual difference might be enlarged when we use photographs of real-life objects as stimuli. This information sampling factor can be investigated by measuring and controlling observers’ gaze or attention in the future. Also, further work is needed to investigate the relationship between individual difference in low-level perception and that in high-level semantic understanding of material classes and perceptual attributes.

In this work, we present the possibility of “True” individual difference in perceiving translucency, which might be the result of each observer’s information sampling, their unique knowledge about physical process of translucency, life experiences, and cultural backgrounds. However, we need further studies to tear apart the “individual difference from data”, which could arise from real differences between individuals, but also from measurement error in the visual judgement task (Mollon et al.,2017). Nevertheless, the individual difference we described in this study can indicate the level of ambiguity of the stimuli and relate to the level of information processing, which is important to be considered in building computational models of material perception.

### The role of image data set

In this study, we include 300 images from various material categories, including fruits, meat, crystals, wax, soap, metal and etc., and do not intentionally balance the number of images within each category. In order to analyze individual observers’ performances across tasks, we need a balance between image categories and the number of trials such that the experiment will be finished in a reasonable time frame. In addition, we also do not control lighting and photography style in the images, which might shift some of the rating results. In general, we believe our image data-set is representative of the real world but it has limitations in both diversity and scale. In the future, we would like to scale up the current experiment with a more balanced image data-set, and to extend the study of individual and group differences across cultural backgrounds, languages, and ages through crowd-sourcing platforms.

In our study, we segment the objects and place them against a uniform background because nearby objects in the background might bias our results such as providing additional cues of object identity and lighting environment. For example, having the background of a kitchen table with other food items might significantly affect how observers categorize the target object as food or non-food. Here, we intentionally exclude the background to allow the observers to focus only on the object itself. In this way, we can isolate the effect of color on material categorization and property estimation. But we acknowledge the lack of background might increase ambiguity of material identity and associated semantic attributes’ judgement. We will systematically investigate how the background context affect translucent perception in future studies.

### Limitation of online study

Due to Covid-19, we held the experiments sessions online, and could not fully control the device used by the participants other than asking everyone to use a laptop with at least 13 inches display and set the monitor to maximum brightness. We recognize using different devices might affect results, but we still find systematic effect of color conditions in our experiments, suggesting the effect of the device isn’t significant.

## Conclusion

Using a diverse data-set of photographs of real objects, we discover that color plays significant roles on translucency perception using three tasks, binary classification, semantic attribute rating, and material categorization. In binary classification and material categorization, converting images to grayscale results in disagreements among observers, therefore, flips the translucency classification label for some images. Removing color also leads to substantial changes in the distribution of semantic attribute ratings for a small proportion of images. At the level of individuals, semantic attribute ratings can be used to predict observers’ own binary classification results in both color and grayscale conditions. In the material classification task, we show that removing color alters observers’ perception of material categories for some images. In particular, observers tend to misjudge images of food as non-food and vice versa in grayscale. Together, our findings reveal a significant but complex role of color on material perception of translucent objects in multiple tasks. On the methodological level, our findings highlight the importance of considering individual difference in material perception.

