## Supplementary Material for "Crystal or Jelly? Effect of Color on the Perception of Translucent Materials with Photographs of Real-world Objects"

**Appendix 1.** The label flipped images in Figure.3 for the results of the Binary Classification task (Experiment 1).
Based on the binary classification result, we label an image as "Translucent" (T) if at least 60% of observers classify it as translucent, as "Opaque" (O) if at least 60% of observers classify it as opaque, as "Unsure" (U) if less than 60% of observers classify it as either translucent or opaque. Using these labels, we find that some images flip from "Translucent" in color condition to "Opaque" in grayscale condition, and vice versa.

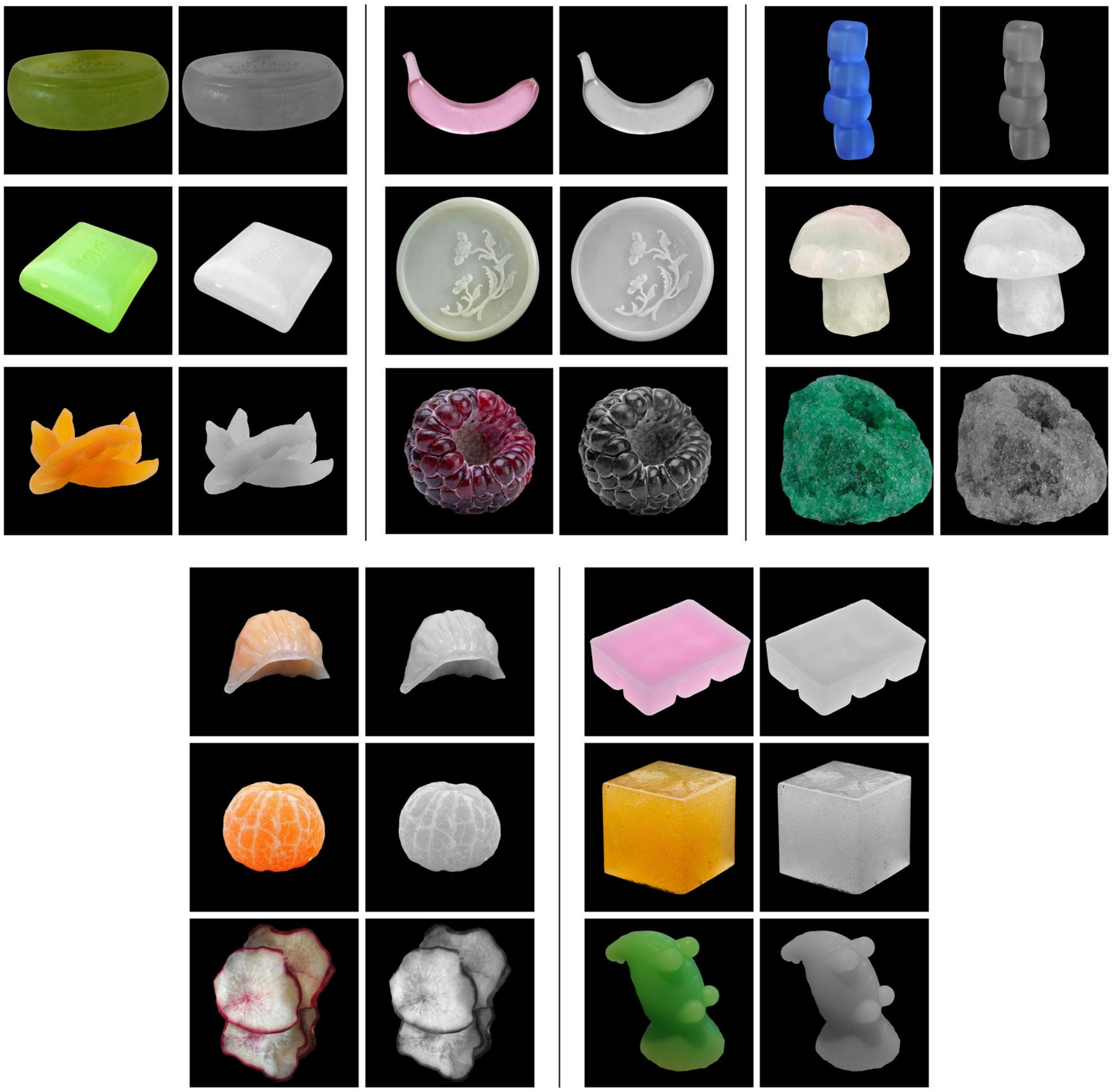

Figure S1.1 The experimental images whose label flips from Translucent to Opaque when they are converted to grayscale by extracting their lightness channel.

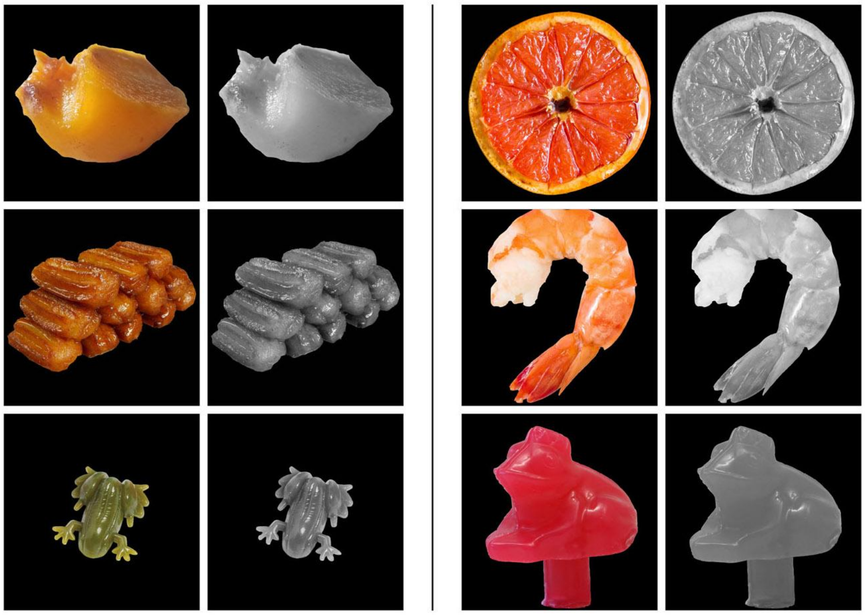

Figure S1.2 The experimental images whose label flips from Opaque to Translucent when they are converted to grayscale by extracting their lightness channel.

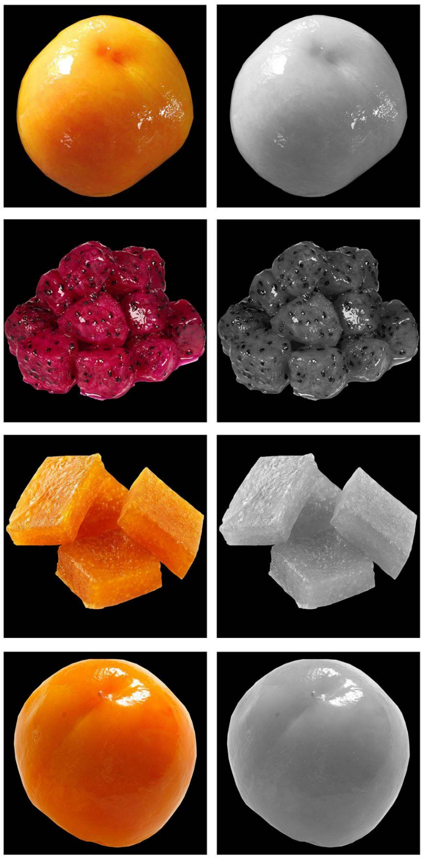

Figure S1.3 The experimental images whose label flips from Unsure to Translucent when they are converted to grayscale by extracting their lightness channel.

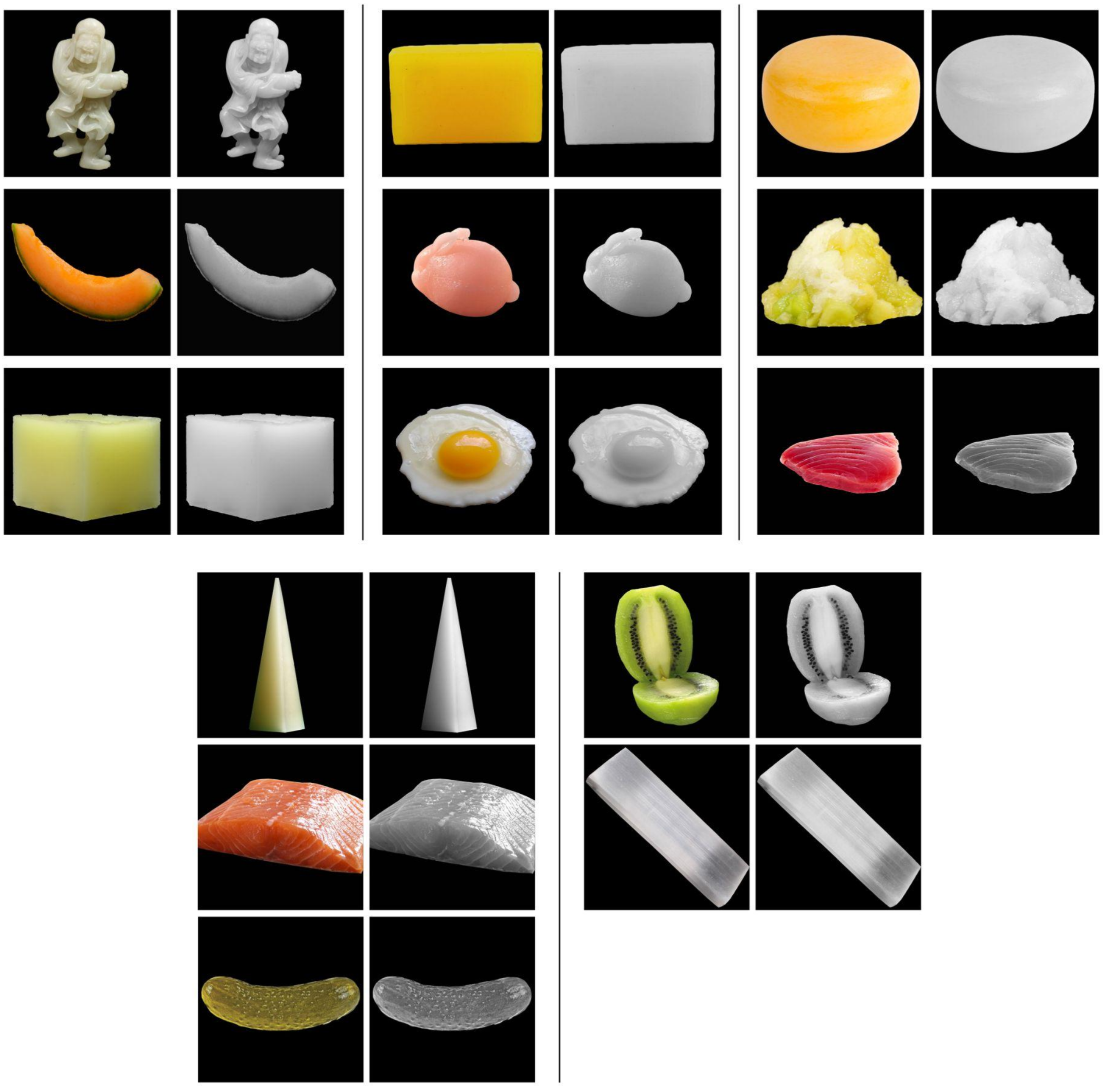

Figure S1.4 The experimental images whose label flips from Unsure to Opaque when they are converted to grayscale by extracting their lightness channel.

**Appendix 2.** Control Experiment: Binary Classification and Material Categorization with Contrast-Preserved Grayscale Images

**Overview**

We conducted an additional experiment with another 20 observers (14 females, 6 males) performing the Binary Classification and Material Categorization tasks using grayscale images with an alternative method, Contrast Preserving Decolorization (Lu et al. (2012)) , (Gray-CP condition).

**Observers**

The participants (12 females and 8 males) have a median age of 19, and all report to have normal color vision. 12 of them used the MacOS system, with 13-inch to 15-inch screens (except one used a 24-inch screen). 8 reported to use the Windows system, with 13-inch to 15.6-inch screens (except one used a 23.8-inch screen).

**Procedure**

The observers first completed the Binary Classification task and then the Material Categorization task. The order of images in each task was identical to that in the grayscale condition (Lightness channel) in the manuscript.

**Results**

Here, we report the results of both Binary Classification and Material Categorization using a new set of grayscale images obtained with the Contrast Preserving method. Figure S2.1(I) shows the percent agreement of binary classification among the 20 observers. The trend of the figure is very similar to Figure 3 (I), suggesting using a contrast-preserving algorithm does not change the trend of the data. Figure S2.1(II) shows the number of images that flipped their classification labels. The trend of this figure is also similar to Figure 3(II) except slightly more number of images flipped their classification label from translucent or unsure to opaque, and from translucent to unsure. Figure S2.1(III) shows examples of images that flipped their classification labels.

Figure S2.2 shows that converting images to grayscale using Contrast Preserving method also leads to more ambiguity in binary translucency classification for some images, and accompanies with higher level of disagreements among observers in comparison to the color condition.
Figure S2.3 shows that observers also erroneously classified ‘food’ as ‘non-food’ and vice versa, and its pattern is similar to that of the grayscale condition (by extracting Lightness channel) illustrated in Figure 12 (I). Figure S2.4 demonstrates that observers are more likely to judge differently in the Gray-CP condition, and are more likely to have higher dissimilarity compared to the color condition.

Overall, we find that results obtained from the grayscale stimuli generated by two conversion methods are similar. Both methods lead to ambiguity in the binary translucency classification task, and higher level of disagreement among observers in both binary classification and material categorization in comparison to the color condition. However, there is a difference in the images whose labels are flipped. Gray-CP condition has more images flipped from “T to U”, “U to O” and “T to O”, and the exact images are different from those in the manuscript.

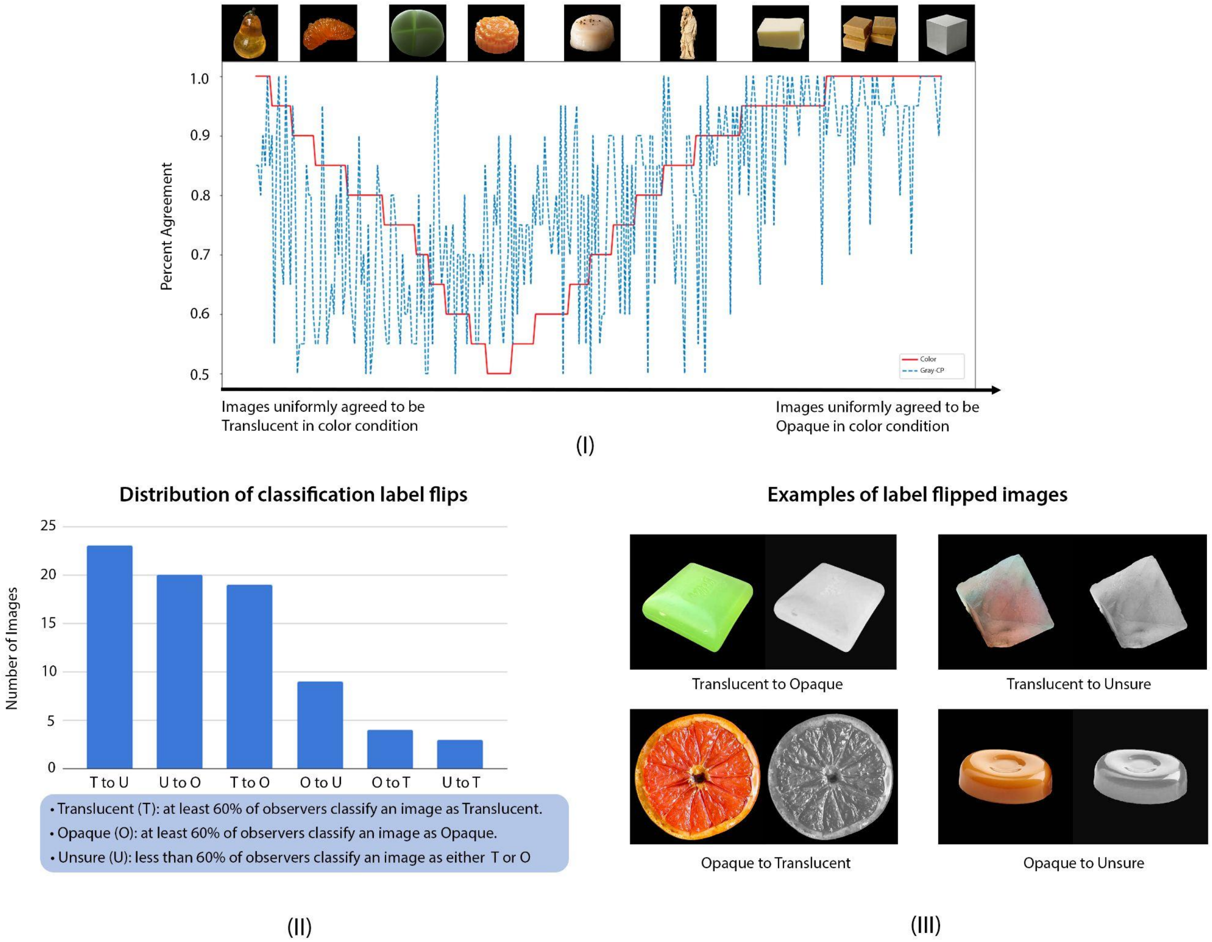


Figure S2.1 Results with the new Gray-CP stimuli. The format of the figure is the same as Figure 3 in the main manuscript except the axis label in panel (II) was ordered as descending order of “Number of Images”.

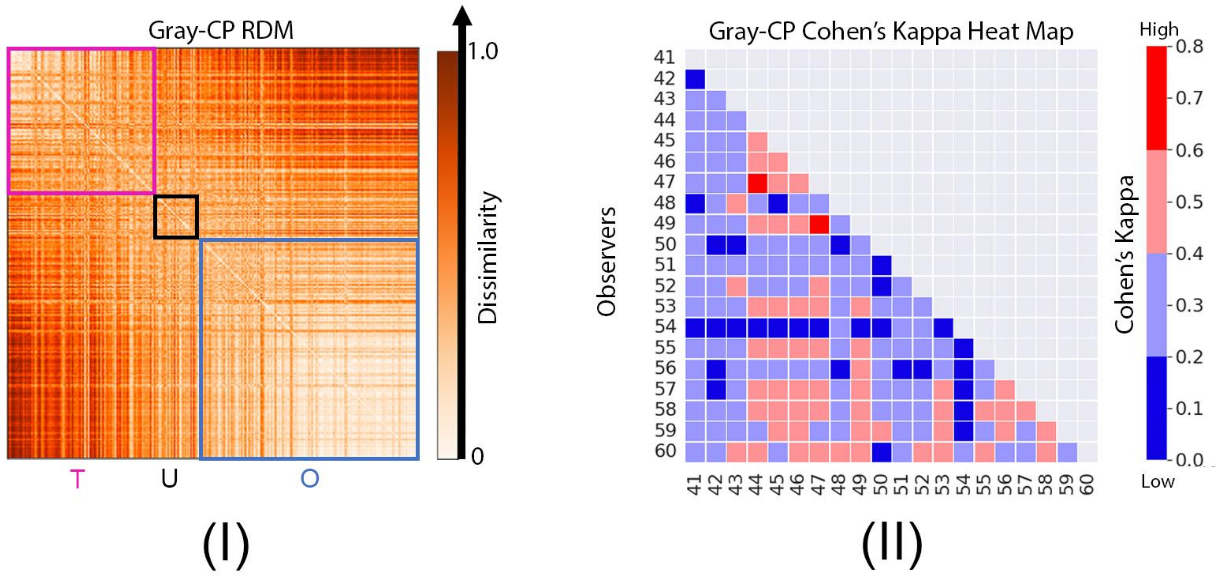


Figure S2.2 Result of the Binary Classification task with Gray-CP condition. The format of the plot is the same as Figure 4 in the main manuscript. Observes in Gray-CP condition are indexed from 41 to 60. (I) RDM of Binary Classification task. We conduct a one-sided Mann-Whitney U test to compare corresponding regions between color and Gray-CP conditions' RDMs and find that all of these regions in Gray-CP RDM have statistically higher dissimilarity in comparison to those in color RDM (p<0.0001 for each pair of regions). (II) Cohen's Kappa heat map of Binary Classification task. Mann Whitney U test comparing the color and Gray-CP conditions shows that observers have lower levels of agreement when the images are shown in the contrast preserved grayscale (p<0.0001).

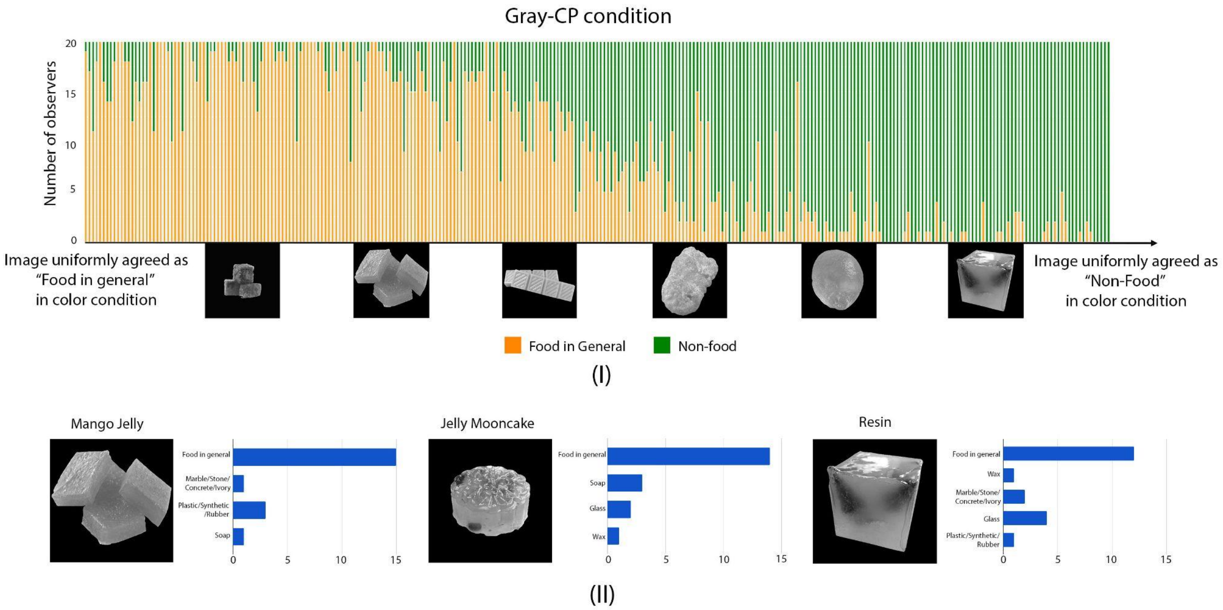


Figure S2.3 Result of the Material Categorization task with Gray-CP condition. The format of the plot is the same as Figure 12 in the manuscript. (I) Trial-by-trial categorization of image as "Food in general" or "Non-food". (II) Examples of images with different material categorization results from observers.

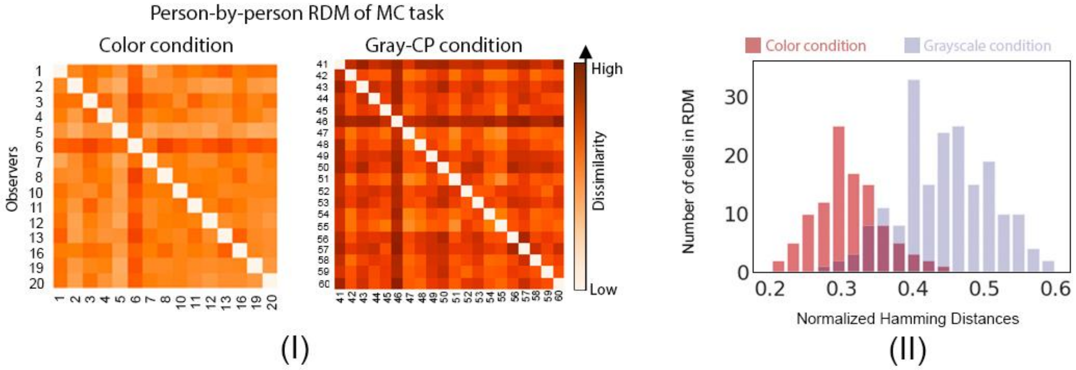


Figure S2.4 Result of the Material Categorization task with Gray-CP condition. The format of the plot is the same as Figure 13 in the manuscript. (I) Person-by-person RDMs of Material Categorization task in color and Gray-CP conditions. We find that the two RDMs are significantly different. Specifically, the Gray-CP condition RDM is more likely to have high dissimilarity values in comparison to the color condition RDM(one-sided Mann-Whitney U test, $p<0.0001$). Unlike Figure 13 in the manuscript, here the range of normalized Hamming distance is from 0 to 0.6. (II) The comparison of distributions of the normalized Hamming distances in color and grayscale condition RDMs.
